# Lipid Accumulation Induced by APOE4 Impairs Microglial Surveillance of Neuronal-Network Activity

**DOI:** 10.1101/2022.03.21.484146

**Authors:** Matheus B. Victor, Noelle Leary, Xochitl Luna, Hiruy S. Meharena, P. Lorenzo Bozzelli, George Samaan, Mitchell H. Murdock, Djuna von Maydell, Audrey H. Effenberger, Oyku Cerit, Hsin-Lan Wen, Liwang Liu, Gwyneth Welch, Maeve Bonner, Li-Huei Tsai

## Abstract

Apolipoprotein E4 (APOE4) is the greatest known genetic risk factor for developing late-onset Alzheimer’s disease and its expression in microglia is associated with pro-inflammatory states. How the interaction of APOE4 microglia with neurons differs from microglia expressing the disease-neutral allele APOE3 is currently unknown. Here, we employ CRISPR-edited induced pluripotent stem cells (iPSCs) to dissect the impact of APOE4 in neuron-microglia communication. Our results reveal that APOE4 induces a distinct metabolic program in microglia that is marked by the accumulation of intracellular neutral lipid stores through impaired lipid catabolism. Importantly, this altered lipid-accumulated state shifts microglia away from homeostatic surveillance and renders APOE4 microglia weakly responsive to neuronal activity. By examining the transcriptional signatures of APOE3 versus APOE4 microglia before and after exposure to neuronal conditioned media, we further established that neuronal soluble cues differentially induce a lipogenic program in APOE4 microglia that exacerbates pro-inflammatory signals. Pharmacological blockade of lipogenesis in APOE4 microglia is sufficient to diminish intracellular lipid accumulation and restore microglial homeostasis. Remarkably, unlike APOE3 microglia that support neuronal network activity, co-culture of APOE4 microglia with neurons disrupts the coordinated activity of neuronal ensembles. We identified that through decreased uptake of extracellular fatty acids and lipoproteins, APOE4 microglia disrupts the net flux of lipids which results in decreased neuronal activity via the potentiation of the lipid-gated K^+^ channel, GIRK3. These findings suggest that neurological diseases that exhibit abnormal neuronal network-level disturbances may in part be triggered by impairment in lipid homeostasis in non-neuronal cells, underscoring a novel therapeutic route to restore circuit function in the diseased brain.

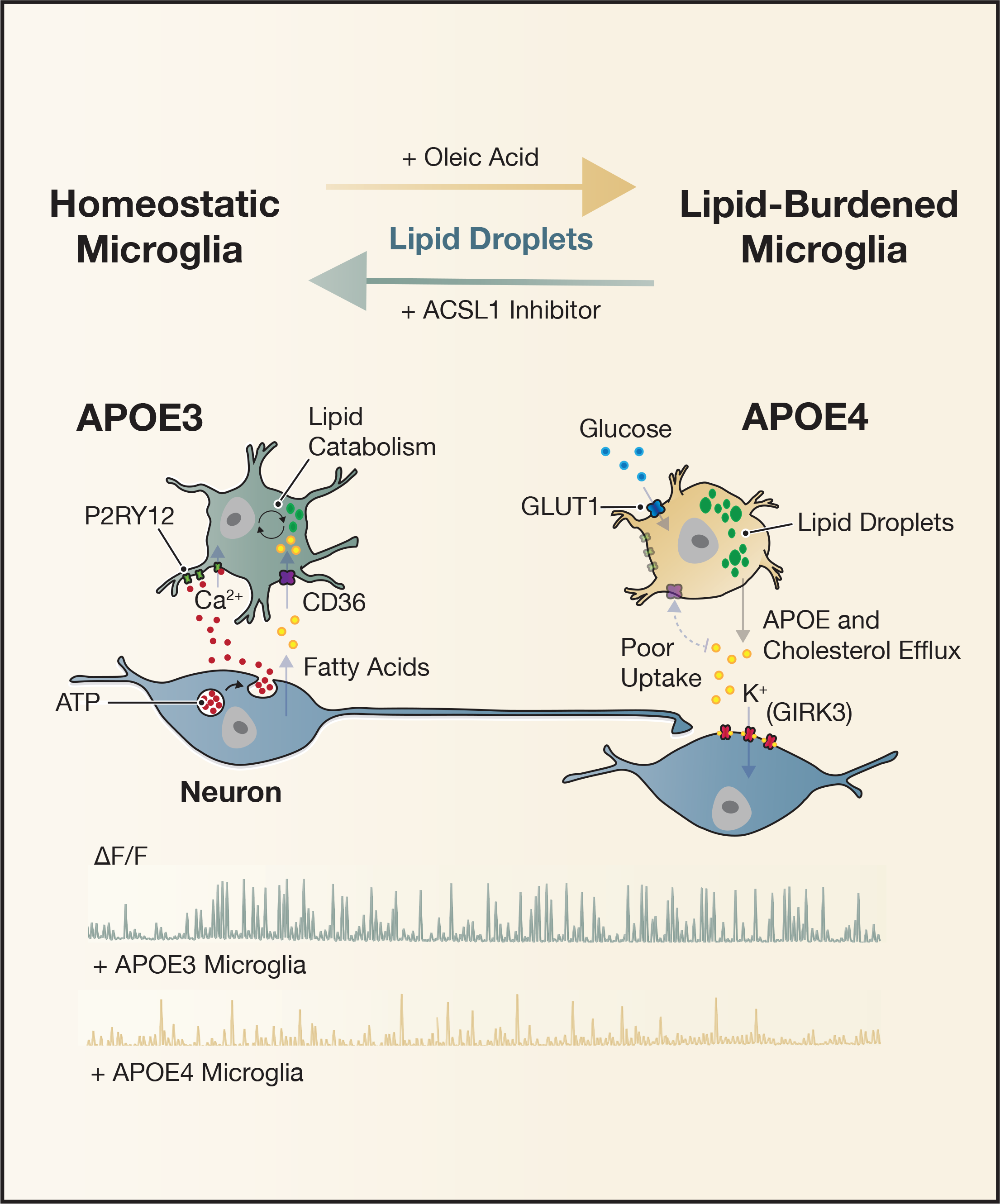

## Introduction

The coordinated regulation of neuronal ensembles is critical for supporting fundamental cognitive processes such as learning and memory (Laubach et al., 2000; Zandvakili and Kohn, 2015). Disturbances to the rhythmic activity of neuronal networks is implicated in many neurological diseases, including Alzheimer’s disease (AD) (Guillon et al., 2017; Koenig et al., 2005). In fact, altered network activity can be detected decades before the onset of clinical symptoms in AD patients (Beason-Held et al., 2013; Sperling et al., 2009), suggesting that it may be an early pathogenic event and a potential therapeutic target to halt or slow down disease progression (Canter et al., 2016; Palop and Mucke, 2016). Recent reports have characterized the close association between microglia and neurons, suggesting that microglia can actively regulate neuronal activity (Badimon et al., 2020; Cheadle et al., 2020; Cserep et al., 2020; Merlini et al., 2021). As such, it has recently been speculated that altered neuronal excitability, which is typical of many neurological diseases, may partly manifest through early deficits in microglial surveillance and regulation of neuronal networks. This is particularly interesting in the context of AD, where a large number of risk variants have been identified in genes that are highly or solely expressed in the brain by non-neuronal cells (Kunkle et al., 2019). Although many glia cell-types are known to modulate and sculpt neuronal circuits, it is unclear how disease-associated genetic drivers enriched in glia, such as single nucleotide polymorphisms (SNPs), impact neuronal network dynamics.

As brain-resident macrophages, microglia are highly reactive to disturbances within the brain microenvironment. This imposes limitations on the viral techniques that are typically employed in systems neuroscience to assess neuron-microglia communication (Maes et al., 2019). While the generation of transgenic mouse models harboring disease-associated alleles circumvents this technical challenge, the rapid pace of human genomic studies far outpaces the low throughput nature of deriving transgenic murine lines. This underscores the need for the development of human-based models to interrogate disease-associated genomic variants in complex multicellular platforms. Furthermore, these systems must be tractable for genome editing with reproducible phenotypic read- outs. Although many studies have employed patient-derived induced pluripotent stem cells (iPSCs) to assess the functional impact of AD-associated genetic variants in cell-types of interest, the impact of risk variants on multicellular interactions have only recently come to prominence (Penney et al., 2020). This is likely a reflection of the advancement of reprogramming technologies that continue to refine multicellular structures, such as brain organoids and spheroids, allowing for more complex physiological interactions to be investigated (Blanchard et al., 2021). Nevertheless, cellular studies utilizing iPSCs to examine the bi-directional communication between neurons and glia with iPSCs are still lacking.

Apolipoprotein E (APOE) is a polymorphic gene with three common alleles, ε2, ε3 and ε4. While APOE ε3 (APOE3) is the most common allele, occurring with 77% frequency in humans and deemed to be neutral with respect to disease, APOE ε4 (APOE4) is the greatest known genetic risk factor for developing late-onset AD (Liu et al., 2013; Yamazaki et al., 2019). The ε2 allele is associated with a decreased risk of developing late-onset AD, and therefore generally referred to as protective, but nevertheless can increase the risk of certain cerebrovascular diseases (Li et al., 2020). APOE3 and APOE4 protein variants differ by a single amino acid change, with a cysteine to arginine modification at position 112 within exon 4 marking the ε4 APOE allele. Within the central nervous system, APOE is predominantly expressed by glia, although neuronal expression of APOE has been reported in stress-induced conditions (Xu et al., 2006; Zalocusky et al., 2021). By leveraging CRISPR/Cas9 genome-editing with iPSC-induced neural cell types, we have previously corrected iPSCs generated from an APOE ε4/4 homozygote AD patient into APOE ε3/3 or induced APOE ε4/4 in a healthy non-demented APOE ε3/3 donor iPSC line, generating pairs of isogenic lines that are genetically identical except for the modification in the APOE allele (Lin et al., 2018). We previously reported that iPS- derived neurons, astrocytes, and microglia that harbored the APOE4 genotype displayed distinct cell type-dependent phenotypes (Lin et al., 2018). Using these CRISPR-edited APOE isogenic lines, we sought to build upon these studies to test the impact of this AD-associated variant on the cellular communication between neurons and microglia using an iPS-based platform.

## Results

### iPSC-derived Microglia-like Cells (iMGLs) Respond to Soluble Factors Secreted by Neurons in an Activity-Dependent Manner

Purinergic signaling is a powerful modulator of microglia chemotaxis, phagocytosis, and pro-inflammatory cytokine production (Davalos et al., 2005; Koizumi et al., 2007; Monif et al., 2009). Within the brain, the purinergic receptor P2RY12 is highly and predominantly expressed by microglia (Cserep et al., 2020), allowing for microglia to rapidly sense extracellular adenosine di- or tri-phosphate (ADP or ATP) secreted by neurons under physiological (i.e. co-released with neurotransmitters) or pathological conditions (i.e. upon infliction of cellular damage) (Calovi et al., 2019). In addition, many soluble neuronal factors maintain microglia in an inactivated surveillance state, such as the CX3CL1-CX3CR1 signaling axis (Finneran and Nash, 2019). Interestingly, microglia also express numerous neurotransmitter receptors, including glutamate receptors such as AMPA, NMDA and mGLURs, suggesting that microglia can sense glutamatergic neuronal communication (Figure 1A) (Szepesi et al., 2018). We sought to determine if microglia derived from iPSCs expressed receptors thought to mediate neuron-microglia communication. We began by generating iMGLs using established protocols that have been characterized to yield cells of similar transcriptional composition to microglia isolated from human brains (Abud et al., 2017; McQuade et al., 2018). After 4 weeks of differentiation from iPS-derived primitive hematopoietic progenitors, iMGLs in culture exhibited ramified morphology, stained positive for microglial specific-markers such as IBA1 and P2RY12, and displayed mature electrophysiological properties typical of *ex vivo* microglia in culture via whole cell patch-clamp analysis (Figure 1B and Supplementary Figure S1A-E). Additionally, immunostaining of iMGLs revealed the expression of canonical ion channels P2X1 and THIK-1, critical regulators of microglial homeostasis (Figure 1B) (Izquierdo et al., 2021; Koizumi et al., 2013; Madry et al., 2018). We also detected expression of the voltage-gated potassium channel KCNE3, and the voltage-gated calcium channel CACNA2D4, in addition to the metabotropic glutamate receptor, GLUR7 (Figure 1B). Collectively these findings suggest that iMGL recapitulate expression patterns of receptors governing surveillance of neuronal activity.

**Figure 1.**
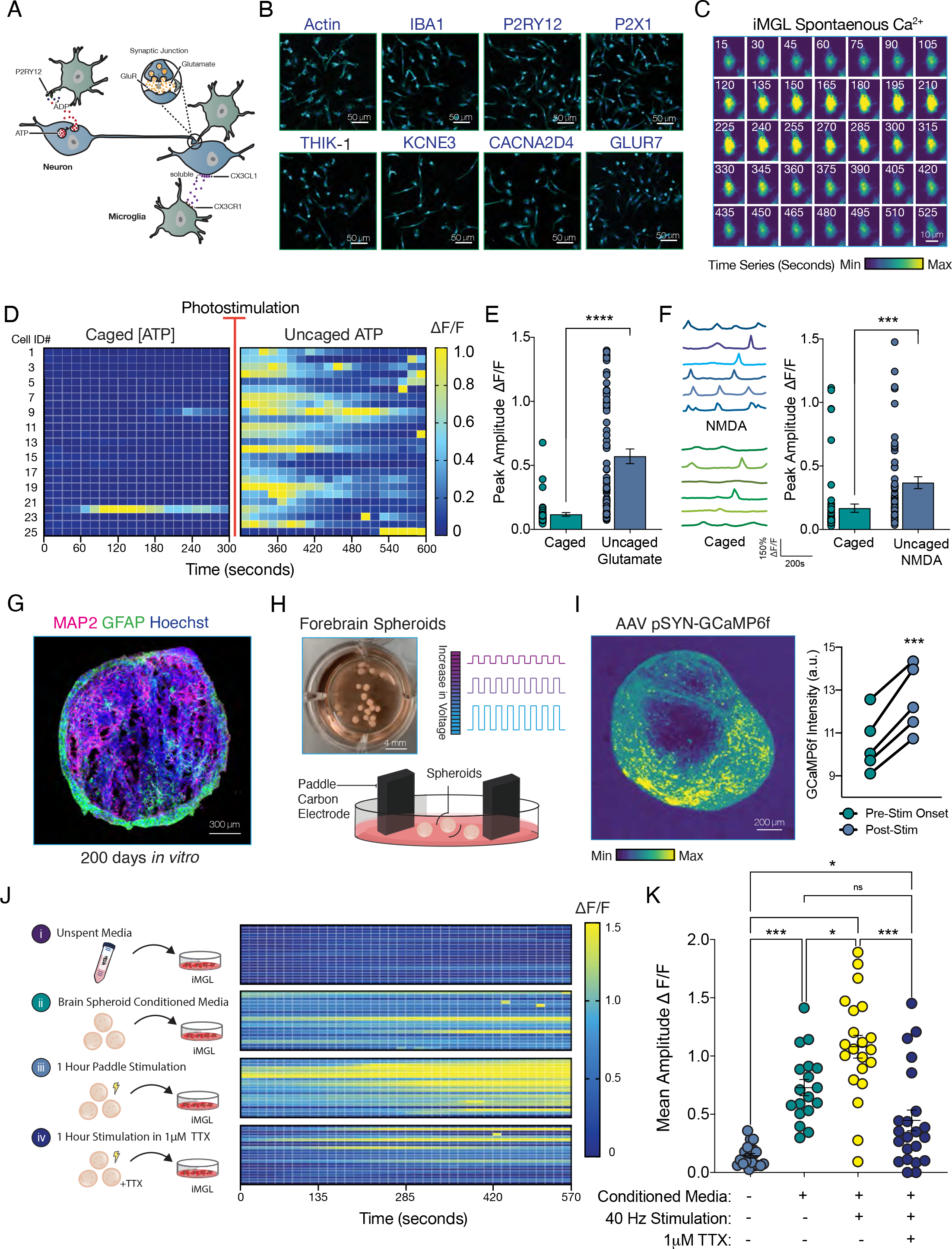
Neuronal Activity Evokes Ca^2+^ Transients in iPSC-derived Microglia-like Cells. (A) Diagram depicting known mechanisms of communication between neurons and microglia. (B) iPSC-derived microglia-like cells (iMGL) express canonical receptors that recapitulate microglia from post-mortem human cortex. (C) Time series of a spontaneous calcium transient observed in monoculture of iMGL with Fluo-4 AM. Changes in signal intensity are displayed from low (blue) to high (yellow) in 15 second bins over 35 frames. (D) iMGL display robust intracellular calcium activation upon detection of ATP; heatmap of iMGL before (left panel) and after (right panel) 405nm photostimulation to uncage 1mM ATP. (E) Change in fluorescence upon uncaging of 5mM glutamate is plotted as peak amplitude for baseline (caged) and post photostimulation (uncaged). n= 64 cells. Paired t-test; *** p-value <0.001 (F) iMGLs can detect extracellular NMDA. Right panel displays representative calcium traces from 6 iMGLs in response to uncaging of 1mM NMDA (upper blue traces). Baseline levels are shown in lower green traces. n= 60 cells. Paired t-test; *** p-value <0.001 (G) Immunostaining of a forebrain spheroid at 5 months for the neuronal marker, MAP2 (Magenta) and the marker for astrocytes, GFAP (Green) with nuclei counterstained with Hoechst (Blue). (H) Forebrain spheroids at 3 months in culture were stimulated to elicit robust neuronal activation using paddle carbon electrodes that dip into media-containing multi-well plates with 15mV electrical pulses for 1 hour. (I) Live-imaging of intact spheroids sparsely labeled with the calcium reporter GCaMP6f expressed under the synapsin promoter (pSYN) upon electrical stimulation. Image of maximum projection across time frames. Calcium imaging prior to stimulation (Pre-Stim) was used to assess baseline levels of calcium transients, and during electrical stimulation onset (Post-Stim). n= 5 spheroids Paired t- test; *** p-value <0.001 (J) Calcium imaging of iMGL in monoculture following 2 hours of incubation in spheroid conditioned media (CM). Data is presented for unspent spheroid media as control (i), CM from intact spheroids (ii), spheroids stimulated for 1 hour (iii), or spheroids stimulated for 1 hour in the presence of the calcium blocker tetrodotoxin (1mM TTX). Full media switches followed the stimulation and media was conditioned for 24 hours before being collected. (K) Peak amplitude of calcium transients from (J) is quantified and plotted. One-Way ANOVA with post-hoc Tukey test. n.s. = not significant; ** p-value <0.01 ; *** p-value <0.001; **** p-value <0.0001. n= 18 to 21 cells in each group.

To better characterize the functional properties of these transmembrane channels, we examined calcium signaling in iMGLs in response to extracellular cues. Receptor-mediated Ca^2+^ signals are a common transduction mechanism in microglia, particularly downstream of ligand-gated calcium-permeable purinergic receptors (Moller, 2002). In addition, the activation of metabotropic receptors is known to trigger the release of intracellular Ca^2+^ stores in microglia, making calcium imaging an attractive tool to examine microglial response to neuronal soluble cues (Farber and Kettenmann, 2006). Given microglia response to viral infections, we reasoned that non-viral mediated approaches to visualize calcium transients may be better suited to avoid evoking microglial activation. Thus, we labeled monocultures of iMGLs with the membrane-permeant calcium indicator, Fluo-4 AM. At baseline, we observed sparse microglial calcium transients, which were characterized by their high amplitude (greater than 1 *Δ*F/F; change in fluorescence over baseline) and prolonged periods (nearly 4 minutes in duration), in line with previous findings of spontaneous calcium dynamics of microglia in awake mice (Umpierre et al., 2020) (Figure 1C). Since iMGLs have been shown to respond in culture to extracellular ATP and ADP in a P2RY12-dependent manner, we next attempted to evoke calcium transients by pharmacologically exposing iMGLs to ATP (Abud et al., 2017; Konttinen et al., 2019). We used a biologically inactive analog of ATP that can be photostimulated with ultraviolet light for rapid activation (Caged ATP Vs. Uncaged ATP) to track the cellular response of the same cell over the course of 10 minutes. As expected, ATP uncaging elicited a robust increase in calcium transients in iMGLs (Figure 1D, Supplemental Movie 1). In addition, we also determined that iMGLs sensed extracellular ATP with whole-cell patch-clamp analysis (Supplementary Figure S1F). Since microglia are known to respond to excitatory neurotransmitters *in vivo (Eyo et al., 2014)*, we sought to determine if calcium transients in iMGLs were elevated upon glutamate exposure. Using a similar drug application strategy to caged-ATP, we found that uncaging NMDA, an amino acid derivative that acts as an NMDA receptor agonist, or glutamate, both increase the amplitude of calcium transients (Figure 1E and Figure 1F). This suggested that iMGLs may sense secreted neuronal soluble factors, such as nucleotides or neurotransmitters.

To test the possibility that iMGLs could respond to a more physiological neuronal stimulus, we next generated forebrain spheroids that were mainly composed of excitatory neurons following previously established protocols (Gordon et al., 2021; Sloan et al., 2018; Yoon et al., 2019). Spheroids are smaller in size than brain organoids (only 1-2mm in diameter) and circumvent the nutrient-poor inner core of large organoids that result in necrotic hotspots. As previously reported, extended culture times of greater than 120 days yield a small, but growing population of GFAP-positive astrocytes amongst MAP2-positive neurons (Figure 1G) (Sloan et al., 2017). Although astrocyte-derived soluble cues have been shown to be critical signaling nodes of the brain microenvironment, we exclusively conducted our studies on spheroids between 60-90 days to restrict our findings to neuronal factors. Since a high level of neuronal activity is associated with a larger number and higher amplitude of microglial calcium transients *in vivo (Umpierre et al., 2020)*, we postulated that pacing spheroids with electrical pulses would stimulate greater release of neuronal soluble factors. We adapted paddle carbon electrodes built onto a tissue culture multi-well plate typically employed over the course of several days to enhance the maturation of iPSC-derived cardiomyocytes (Figure 1H) (Ronaldson-Bouchard et al., 2018). To optimize the pacing of spheroids, we infected 3D cultures with Adeno-Associated Virus (AAV) carrying the genetic calcium indicator GcaMP6f under the neuron-restricted synapsin promoter (AAV pSYN-GcaMP6f). Penetration of AAV into these cultures successfully labeled a subset of surface level neurons (Figure 1I). Stimulation with paddle electrodes evoked a robust and sustained neuronal response as quantified by the change in pre-stimulation vs. the post-stimulation GCaMP fluorescence (Figure 1I and Supplemental Movie 2).

To determine if iMGLs responded to neuronal secreted factors in an activity-dependent manner, we performed media carry-over experiments in four experimental groups (Figure 1J). Unspent neuronal media evoked no response in iMGLs after 2 hours of incubation, while neuronal media conditioned for 24 hours with forebrain spheroids significantly increased the number of calcium transients, as indicated by the elevated mean amplitude in *Δ*F/F (Figure 1J,K). When we treated iMGL with conditioned media from spheroids that had been stimulated with paddle electrodes for 1 hour, we observed increased calcium transients of iMGLs (Figure 1J,K). Importantly, when the stimulation was performed in the presence of tetrodotoxin (TTX), a potent inhibitor of neuronal activity, the increase in iMGL calcium transients mediated by stimulated spheroid media was reduced to levels of non-stimulated spheroid media (Figure 1K). To avoid carrying-over TTX to the microglia, stimulation was followed by three washes in warm media and a complete media switch free of TTX that was allowed to condition for 24 hours. Non TTX-treated stimulated controls were handled identically to control for drug washout manipulation. Collectively, these experiments demonstrate that iMGLs can sense neuronal soluble cues in an activity-dependent manner and establish an experimental platform to interrogate the impact of AD-associated risk factors on neuron-microglia cellular communication.

### Modeling Neuron-Microglia Communication with CRISPR-edited APOE3 and APOE4 iMGLs

We envisioned that combinatorial experiments mixing and matching forebrain spheroids and iMGLs derived from CRISPR-edited isogenic iPSC lines harboring either APOE3 or APOE4 alleles could be a powerful approach to determine the functional impact of APOE4 on neuron-microglia communication. We began by generating iMGLs from APOE3 and APOE4 iPSCs and characterizing their calcium transients with Fluo-4 AM (Figure 2A and Figure 2B). At baseline, we did not observe any differences in the levels of spontaneous calcium transients between genotypes (Figure 2C). To determine if iMGLs responded differently to application of neuronal soluble factors, we applied APOE3 spheroid conditioned media (CM) to monocultures of APOE3 or APOE4 iMGLs and measured calcium transients over the same recording window. Interestingly, neuronal conditioned media evoked fewer calcium transients in APOE4 than APOE3 iGMLs (Figure 2D,E). In addition, upon ATP uncaging, APOE4 iMGLs displayed a blunted response compared to APOE3 iMGL controls (Figure 2F). iMGLs derived from a distinct donor parental line showed similar responses suggesting that this phenotype is independent of isogenic cell line derivation (Figure 2G). Although the blunted response observed in APOE4 iMGLs exposed to neuronal CM could indicate a deficit in a number of signaling systems, ATP uncaging experiments indicate that APOE4 iMGLs are particularly impaired in purinergic signaling. These results suggest that APOE4 iMGLs are weakly attuned to neuronal activity. Moreover, given that downregulation of purinergic receptors, namely P2RY12, is associated with microglial activation status, we reasoned that homeostatic surveillance state is shifted in APOE4 iMGLs.

**Figure 2.**
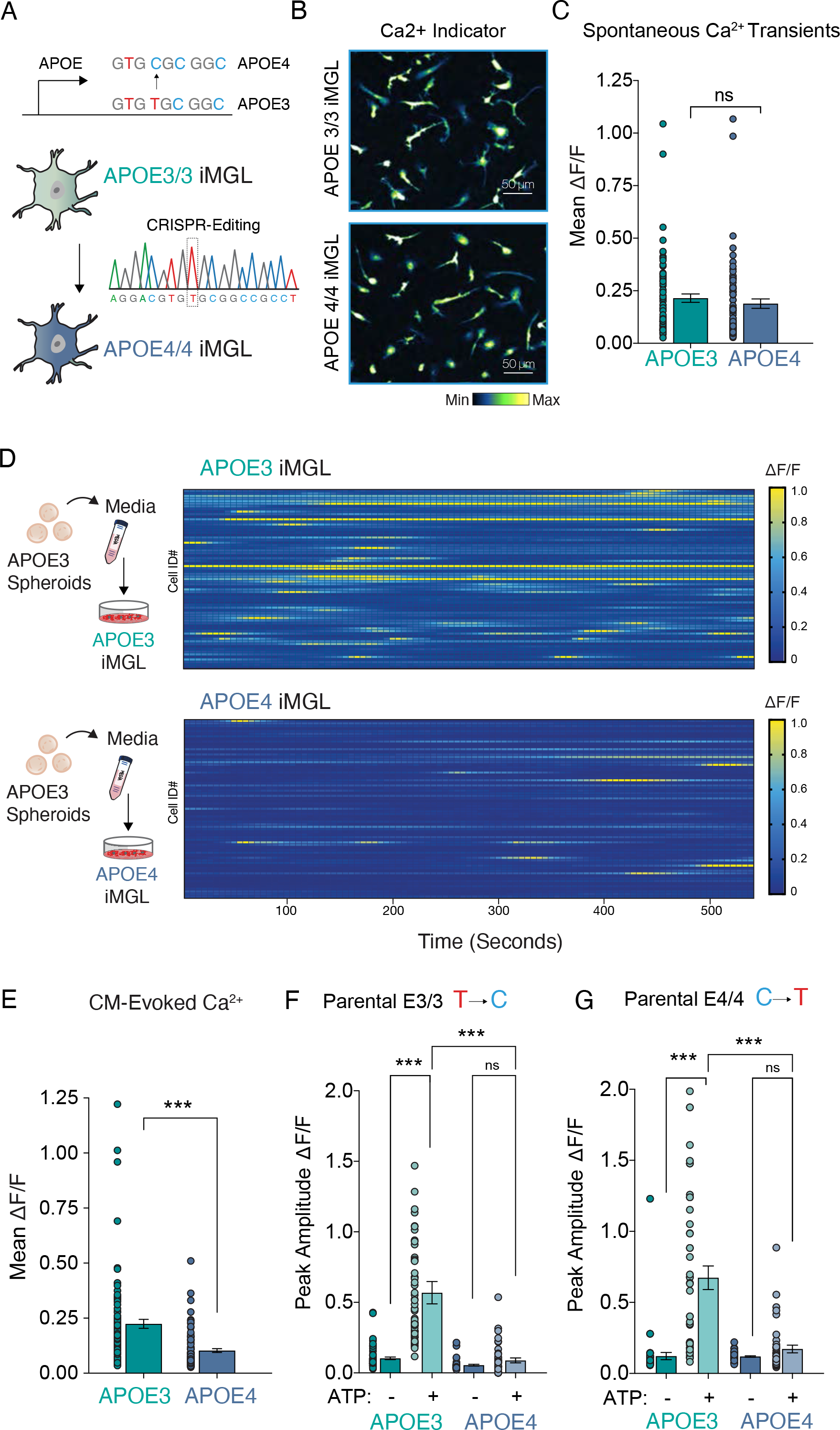
APOE4 iMGL are Weakly Attuned to Neuronal Activity. (A) Diagram of CRISPR- generated isogenic APOE E3/3 (APOE3) and E4/4 (APOE4) iPSCs that were subsequently differentiated into iMGLs. (B) Maximum intensity projection of calcium imaging of APOE3 and APOE4 iMGLs with Fluo-4 AM; fluorescence intensity is shown in gray scale. (C) Number of spontaneous calcium transients did not significantly differ between genotypes, quantified as average change in fluorescence. Unpaired t-test; ns=not significant ; n= 70-80 cells in each group. (D) Heatmap from isogenic pairs incubated with conditioned media (CM) from stimulated APOE3 forebrain spheroids for 2 hours, and (E) quantified as mean change in fluorescent intensity. Unpaired t-test; **** p-value <0.0001; n= 90 cells in each group. (F-G) ATP uncaging evokes robust calcium transients in APOE3 iMGLs, while APOE4 iMGLs show a blunted response independent of isogenic line derivation, (F) APOE3 parental line; n=39-45 cells per group and in (G) APOE4 parental line; n= 41-65. ANOVA with post-hoc Tukey test. n.s. = not significant; ** p- value <0.01; *** p-value <0.001; **** p-value <0.0001.

### Neuronal Conditioned Media Evokes Distinct Transcriptional Responses in APOE3 Vs. APOE4 iMGLs

Using an alternative strategy for deriving microglia-like cells from iPSCs, we have previously reported that transcriptional differences between APOE4 and APOE3 microglia reflect an enhanced inflammatory state (Lin et al., 2018). These data were collected in the absence of neuronal exposure, and therefore to dissect the mechanisms whereby APOE4 reduces microglial surveillance of neuronal activity, we probed the transcriptional profile of these cells at baseline and in response to spheroid conditioned media. After 4 weeks in culture, APOE3 or APOE4 iMGLs pre-conditioned in unspent neuronal media for 24hrs were further incubated with spheroid conditioned neuronal media for 2 hours. Cells were lysed and harvested for RNA extraction, library preparation and bulk sequencing. Biological triplicates were analyzed for 4 groups: APOE3 vs APOE4 iMGLs with or without exposure to APOE3 spheroid conditioned media (+CM) (Figure 3A). Principal component analysis (PCA) revealed that samples clustered most by genotype and conditioned media exposure (Figure 3A). Comparison of APOE3 to APOE4 iMGLs revealed 4,167 differentially expressed genes (DEGs) (APOE3 vs APOE4 iMGLs; False Discovery Rate (FDR) corrected p-value <0.05) (Supplementary Figure S2A).

**Figure 3.**
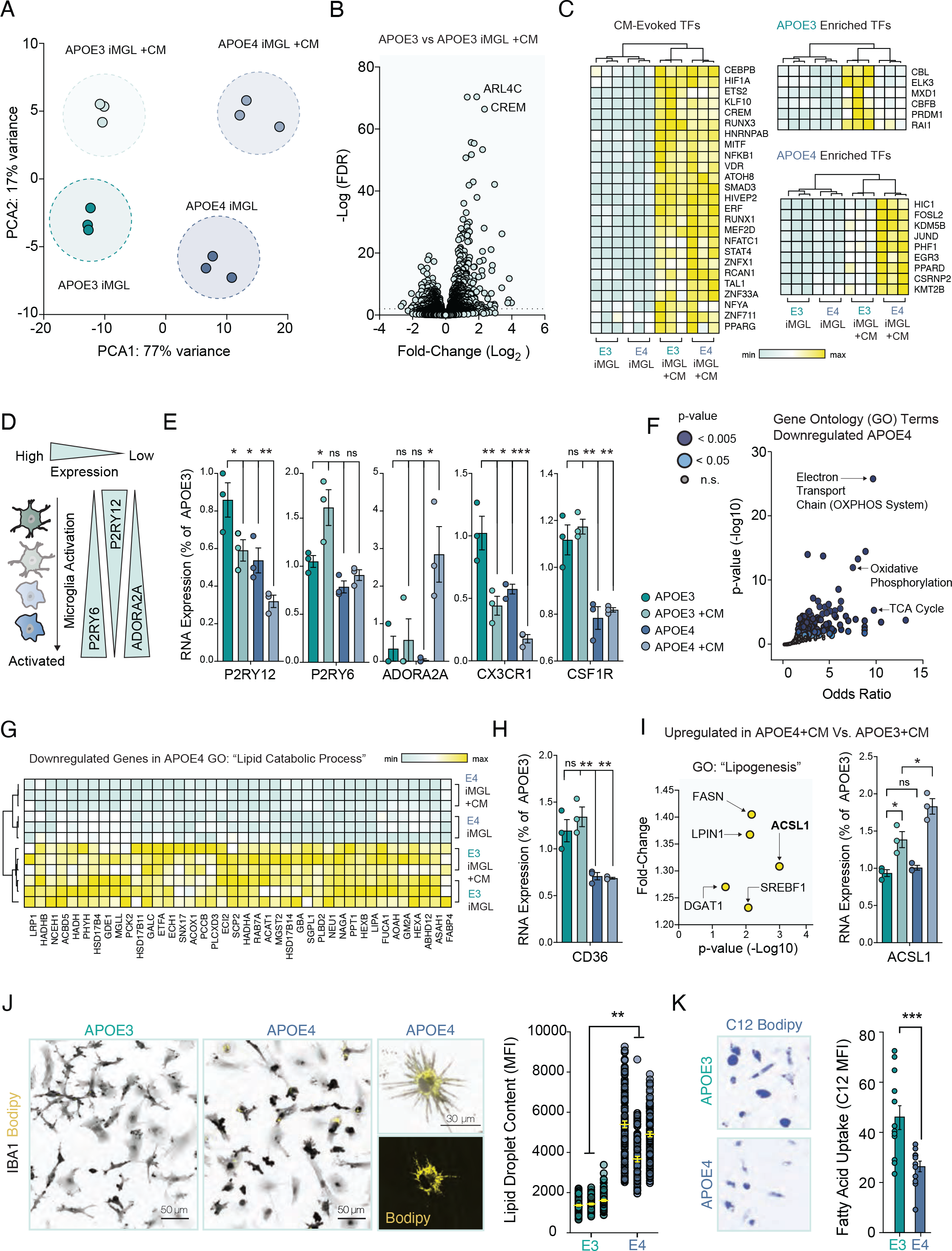
APOE4 Shifts iMGL into a Metabolic Distinct Cell State That is Marked by Impaired Lipid Catabolism. Transcriptional analysis (RNA-seq) of APOE3 and APOE4 isogenic iMGLs exposed to APOE3 spheroid conditioned media (CM) for 2 hours. (A) Principal Component Analysis (PCA) of biological triplicates for 4 groups (APOE3 iMGL +/- CM and APOE4 iMGL +/- CM). (B) Volcano plot of differentially expressed genes (DEGs) evoked by spheroid conditioned media in APOE3 iMGLs (APOE3 vs. APOE3 iMGL + CM). Dotted line indicates FDR cut-off of 0.01. Relative gene expression is shown from low (green) to high (yellow). (C) Heatmap of differentially expressed transcription factors (TFs) evoked by CM. Differentially expressed TFs unique to APOE3 or APOE4 are shown separately but derived from the same dataset. (D) Diagram depicting microglial activation and associated changes in purinergic signaling. (E) Normalized read counts from RNA-seq DEGs are shown as histograms for P2RY12, P2RY6, ADORA2A, CX3CR1 and CSF1R. ANOVA with post-hoc Tukey test. n.s. = not significant; * p-value <0.05; ** p-value <0.01; *** p- value <0.001. (F) Gene ontology (GO) pathways for down-regulated DEGs in APOE4. (G) Heatmap for DEGs associated with lipid catabolism. (H) Levels of CD36, a fatty-acid transporter, shown as a histogram from normalized read counts from RNA-seq DEGs. ANOVA with post-hoc Tukey test. n.s. = not significant; ** p-value <0.01. (I) DEGs associated with lipogenesis, upregulated in APOE4 +CM. Histogram for ACSL1 levels across all groups. ANOVA with post-hoc Tukey test. n.s. = not significant; * p-value <0.05. (J) APOE4 iMGLs accumulates intracellular neutral lipid compared to APOE3 iMGL. BODIPY staining (yellow) in APOE3 versus APOE4 iMGLs (counterstained with IBA1 in gray) shows strong accumulation of lipid droplets in APOE4 iMGLs. n= 73-107 cells per group in 3 separate experiments. Averages from the 3 groups were used for unpaired t-test; ** p-value <0.01. (K) 24 hours of incubation with the green fluorescent fatty acid, C12 Bodipy (shown in blue), reveals decrease uptake by APOE4 iMGLs. Unpaired t-test; **** p-value <0.0001; n= Averages from 12 separate replicates in each group with 16-33 cells quantified per replicate.

To understand how spheroid conditioned media affects iMGL transcription at baseline, we first performed differential analysis comparing APOE3 to APOE3+CM. We identified 604 downregulated and 884 upregulated DEGs in APOE3+CM iMGLs (Figure 3B). Gene ontology (GO) analysis of CM-evoked transcripts in APOE3 iMGLs revealed a strong signature of secondary signaling cascades including cAMP signaling (FDR 7.19x10^-3^), Phospholipase D signaling (FDR 9.14x10^-4^), MAPK signaling (FDR 1.33x10^-4^) and the regulation of actin cytoskeleton (FDR 1.29x10^-2^). (Supplementary Figure S2D and Supplemental Table 1). In fact, one of the most highly enriched genes induced by conditioned media was the cAMP response element modulator, CREM (APOE3 vs. CM; FDR 4.08x10^-67^). Increased cAMP-mediated signal transduction in microglia is associated with the rapid generation of actin-dependent filopodia which allows for fast nanoscale surveillance within discrete regions of the brain parenchyma (Bernier et al., 2019). In contrast, APOE4 iMGL exposure to spheroid conditioned media evoked a larger transcriptional response with 1,305 down and 1,702 upregulated DEGs. APOE4+CM iMGLs were significantly enriched for HIF-1 signaling (FDR 2.26x10^-2^), JAK-STAT signaling (FDR 2.15x10^-2^), Cytokine-cytokine receptor interaction (FDR 2.59x10^-3^), and Phagosome (FDR 1.54x10^-3^) (Supplementary Figure S2D and Supplemental Table 2) suggesting a strong pro-inflammatory response. Similar to APOE3 iMGLs however, spheroid conditioned media also evoked upregulation of cAMP-mediated transcripts in APOE4 iMGLs, including CREM, although to a lesser extent (Supplemental Table 2). In fact, we observed decreased induction of several targets of intracellular Ca^2+^ signaling in APOE4 iMGLs+CM in direct comparison to APOE3 iMGLs+CM, including calmodulins (e.g. CALM2 and CALM3), mitogen-activated protein kinases (e.g. MAPK1 and MAPK9), and calcium/calmodulin-dependent protein kinases (e.g. CAMK2D and CAMK1) (Supplementary Figure S2F), congruent with our observation of decreased calcium transients in APOE4 iMGLs in spheroid conditioned media.

Our initial differential expression analysis revealed that both APOE3 and APOE4 iMGLs upregulate Ca2+ signaling pathways in response to spheroid conditioned media, although this pathway enrichment is muted in APOE4 iMGLs. Additionally, APOE4 iMGLs seem to activate more inflammatory pathways in response to spheroid conditioned media than APOE3 iMGLs. To further dissect the differences between APOE3 and APOE4 iMGLs in response to conditioned media, we directly compared their transcriptional profiles (APOE3+CM vs. APOE4+CM). Emerging genetic mechanisms have linked the mobilization of intracellular Ca^2+^ with downstream lipid signaling, particularly through the induction of lipid secondary messengers such as phosphatidylinositol 3-kinases (PI3Ks) (Vanhaesebroeck et al., 2012). In microglia, PI3K-AKT signaling regulates many cellular functions, including the production of cytokines in response to pro-inflammatory stimuli (Cianciulli et al., 2020). Although we detected moderate enrichment of the PI3K-AKT signaling pathway in APOE3 iMGLs exposed to spheroid conditioned media (APOE3 vs. APOE3+CM) (Supplemental Table 1), we observed a comparable increase in the induction levels of AKT serine/threonine kinase 1 (AKT1) in APOE4 iMGLs +CM in comparison to APOE3 iMGLs +CM (Supplementary Figure S2F). AKT hyperactivation has been recently reported in microglia of AD patients, and pharmacological inhibition of AKT in murine microglia was found to reverse inflammatory signatures (Sayed et al., 2021). Also of note regarding the relationship between calcium influx and lipid signaling is the upregulation of ARL4C (Also known as ARL7), the most overall enriched DEG we identified in CM-evoked transcripts in APOE3 (Log2 fold-change = 1.87, FDR 4.05x10^-71^) and even more so in APOE4 (Log2 fold-change = 2.43, FDR 6.20x10^-191^) (Figure 3B). The ADP-ribosylation factor-like 4C or ARL4C, is a direct target induced by the activation of liver X receptor (LXR) and has been shown to transport cholesterol to the membrane for ABCA1-associated removal in macrophages (Hong et al., 2011). Therefore, our initial analysis implicates lipid metabolic and inflammatory gene programs in the response of iMGLs to conditioned media.

To dissect the regulatory landscape that governs the distinct lipid metabolic and inflammatory transcriptional response by APOE4 iMGLs to spheroid conditioned media, we identified transcription factors (TFs) significantly enriched in either APOE3 iMGLs, APOE4 iMGLs, or both genotypes upon exposure to spheroid conditioned media (Figure 3C). Commonly evoked TFs included several master regulators of inflammation, including the nuclear factor kappa B subunit 1 (NFKB1), and the signal transducer and activator of transcription 4 (STAT4). We observed a distinct set of inducible TFs that bifurcated on the known role to mitigate or exacerbate inflammatory processes in a genotype-dependent manner (Figure 3C). For instance, neuronal media induced expression in APOE3 iMGLs of the proto-oncogene CBL which has been shown to repress pro-inflammatory activation pathways in immune cells (Lu et al., 2021; Shamim et al., 2007; Zhang et al., 2003), and also PRDM1 (Positive Regulatory Domain 1, also known as BLIMP1) which was identified as a repressor of interferon gene expression (Keller and Maniatis, 1991), with depletion of PRDM1 being associated with aberrant and exacerbated activation of inflammatory reactions (Fairfax et al., 2007; Kim et al., 2011). In addition, the transcriptional repressor ELK3, which is also uniquely induced in APOE3 iMGLs by neuronal conditioned media, has been shown to directly repress the expression of pro-inflammatory nitric-oxide synthase in macrophages (Chen et al., 2003), with its transcript being downregulated in a dose-dependent manner in response to bacterial lipopolysaccharide (LPS) (Tsoyi et al., 2015). The induction of these inflammatory repressors by APOE3 iMGLs (i.e. CBL, PRDM1 and ELK3) in response to spheroid conditioned media may act as an immune checkpoint to mitigate downstream inflammatory responses despite the induction of immune master regulators. Failure to evoke these inducible-TFs suggests that this immune checkpoint is left unchecked in APOE4 iMGLs, perhaps leading to overactivation of downstream immune effectors. To this point, neuronal inducible-TFs unique to APOE4 iMGLs have been linked with promoting inflammation (e.g. HIC1 and FOSL2) (Burrows et al., 2017; Renoux et al., 2020) or to be induced in macrophages by pro-inflammatory stimuli (e.g. CSRNP2, JUND and EGR3) (Eichelbaum and Krijgsveld, 2014). Although our data suggests that APOE3 and APOE4 iMGLs responded similarly to changes in their microenviroment (i.e. acute exposure to neuronal soluble cues), we observed a dichotomy in the execution of inflammatory responses between APOE4 and APOE3 iMGLs.

Microglial activation is defined by dramatic changes to cell morphology and to purinergic signaling (Figure 3D) (Koizumi et al., 2013). Since our assays indicated that the capacity of APOE4 iMGLs to sense ATP or broader neuronal-secreted soluble factors were decreased, we next examined the levels of P2RY12 across APOE3 and APOE4 iMGLs. APOE4 iMGLs expressed significantly lower levels of P2RY12 than APOE3 iMGLs (Figure 3E). While exposure to spheroid conditioned media for 2 hours significantly decreased levels of P2RY12 in both genotypes, P2RY6 expression levels increased only in APOE3 iMGLs. Upregulation of P2RY6 in microglia is associated with a hypervigilant microglial activation state that acts as a primer for the phagocytosis of dead cells or debris (Koizumi et al., 2013). Interestingly, despite these deficits in purinergic receptor expression, APOE4 iMGLs dramatically increased the expression levels of adenosine receptor A2A (encoded by the gene ADORA2A) (Figure 3E). In the brain, ATP can be rapidly hydrolyzed into adenosine which is a potent activator of microglia pro-inflammatory phenotype (Colella et al., 2018). Upregulation of A2A receptor is associated with the shift from ramified to amoeboid microglial morphology during brain inflammation (Orr et al., 2009). In agreement with this observation, we detected a reduction in multiple homeostatic microglia genes, including CX3CR1 and CSF1R (Figure 3E) which have been reported to be downregulated in activated microglia (Krasemann et al., 2017). While our transcriptional profiling provided molecular insight into the decreased surveillance of APOE4 iMGLs to neuronal soluble cues, we next sought to identify the cellular mechanism by which APOE4 genotype shifts microglial status away from homeostatic surveillance.

### Microglial Energetics and Lipid Processing are altered in APOE4 iMGLs

At baseline and upon spheroid conditioned media, we observed a dramatic decrease in DEGs relating to mitochondrial oxidative phosphorylation (OXPHOS) in APOE4 iMGLs (Figure 3F and Supplementary Figure S2G). Deficits in mitochondrial metabolism that yield a low energy cellular state have been previously reported in APOE4 human subjects, as well as in mice and iPSC-derived glia harboring APOE4 alleles (Farmer et al., 2021; Konttinen et al., 2019; Orr et al., 2019; Qi et al., 2021). Furthermore, pro-inflammatory stimuli are known to induce a metabolic switch in microglia from OXPHOS to glycolysis, modifying the rate of fatty acid catabolism for the production of ATP by instead relying on the consumption of glucose (Lauro and Limatola, 2020). In alignment with this metabolic reprogramming observed in pro-inflammatory microglia, we observed a small but significant upregulation of the glucose transporter GLUT3 in APOE4 iMGLs (Log2 fold-change = 0.37, FDR 0.0315), while GLUT1 dramatically increased in expression (Log2 fold-change = 6.01, FDR 1.67x10^-23^) relative to APOE3 iMGLs (Supplementary Figure S2H). The upregulation of GLUT1 in pro-inflammatory microglia increases glucose uptake and promotes glycolysis (Wang et al., 2019). Moreover, through our transcriptional analysis we observed that HIF-1 signaling is enriched in APOE4 iMGLs. HIF-1*α* is a master transcriptional regulator of glycolysis, and is induced by AKT through phosphorylation of mammalian target of rapamycin (mTOR) (Cheng et al., 2014). Metabolic reprogramming of microglia from OXPHOS to glycolysis by pro-inflammatory stimuli is dependent on the AKT-mTOR-HIF-1*α* pathway (Baik et al., 2019). These results suggest that APOE4 expression in microglia induces a metabolic reprogramming in energy production that is associated with a pro-inflammatory state.

Mitochondrial oxidation of free fatty acids is a critical mechanism by which lipids are broken-down as energy substrates. Energy depletion and impairment of fatty acid oxidation has been associated with intracellular lipid accumulation in activated microglia (Loving and Bruce, 2020). As such, concurrent with downregulation of genes related to OXPHOS, we also observe a dramatic decrease in DEGs related to lipid catabolic processes in APOE4 iMGLs in relation to APOE3, independent of conditioned media application (Figure 3G). Moreover, we also detected a significant downregulation of the membrane fatty-acid transporter CD36 (Also known as FAT) in APOE4 iMGLs in comparison to APOE3 iMGLs (Figure 3H). Defective fatty acid uptake has been previously observed in APOE4 astrocytes (Qi et al., 2021), and CD36 has been shown to be downregulated in AD-associated proinflammatory microglia (Dobri et al., 2021). Since the uptake of fatty acids are associated with its metabolic demand to fuel lipid oxidation (Bruce et al., 2018; Mashek and Coleman, 2006), our transcriptional profiling reveals a molecular program that may lead to the accumulation of lipids via disrupted mitochondrial function. Interestingly, upon exposure to neuronal conditioned media we further detected the differential upregulation of a subset of genes in APOE4 iMGLs involved in the *de novo* production of lipids or in the regulation of its cellular storage, with the Acyl-Coa synthethase ACSL1, showing the most robust enrichment (Figure 3I). Upregulation of ACSL1 has been previously reported in the postmortem human brain of APOE4 carriers, and to be similarly enriched in iPSC-derived astrocytes harboring the APOE4 allele by our own research group (Sienski et al., 2021). To confirm our RNA-seq analysis, we next determined the abundance of intracellular lipids in APOE4 iMGLs, as well as their ability to buffer, or uptake, extracellular free fatty acids. Indeed, staining for intracellular neutral lipid stores known as lipid droplets with the fluorescent dye BODIPY (counter stained with the microglia-specific marker IBA1) reveals significantly greater lipid droplet content in APOE4 iMGLs in comparison to APOE3 iMGLs (Figure 3J). This is in alignment with our previous effort to quantify lipid droplets in APOE4 bearing iMGLs (Sienski et al., 2021). In addition, incubation of iMGLs with the green-fluorescent fatty-acid C12 BODIPY (C12 BODIPY) reveals decreased uptake by APOE4 iMGLs (Figure 3K), indicative of a functional deficit in fatty acid uptake and mirroring our RNA-seq analysis of decreased CD36 expression in APOE4 iMGLs.

In addition to fueling lipid oxidation, fatty acid uptake by glia is particularly important in preventing neurodegeneration. Neurons have minimal capacity to catabolize fatty acids or store lipids and therefore rely on glial mitochondrial oxidation for lipid consumption by transferring fatty acids via APOE-associated particles (Ioannou et al., 2019). Interestingly, the metabolic coupling of neurons and astrocytes is disrupted by APOE4 leading to impaired neuronal synaptic maturation (Qi et al., 2021). Although metabolic coupling with neurons have been predominantly studied in the context of astrocytes, microglia and astrocytes have both been shown to accumulate lipid droplets *in vivo* upon neuronal excitotoxic injury (Ioannou et al., 2019). In addition, accumulation of lipid droplets in microglia have been reported to represent a dysfunctional and pro-inflammatory state in the aging brain (Marschallinger et al., 2020). Yet, the functional repercussions of a lipid burdened microglial state to the activity of neuronal circuits remains unknown.

### APOE4 Microglia Impair the Highly Coordinated Neuronal Activity of APOE3 Spheroids

Although co-cultures of iPS-derived microglia with neurons have been reported to accelerate the emergence of synchronized neuronal network activity (Popova et al., 2021), it remains unclear if disease-associated genetic risk variants expressed by microglia disrupt this process. To assess how APOE4 iMGLs impact the activity of neurons, we began by dissociating spheroids grown in 3D after 60 days into a single-cell suspension and plating these cultures onto coverslips. We reasoned that a 2D culture would give us higher tractability, allowing us to quantify neuronal activity and monitor microglial health in mixed cultures. After 4 weeks in culture, we found that dissociated spheroids displayed mature neuronal morphology with extensive neurite networks in 2D and were free of GFAP-positive cells (Supplementary Figure S3A). Dissociated spheroids were efficiently infected by AAV virus as shown by the expression of EGFP under the neuronal specific promotor Synapsin (AAV pSynapsin-EGFP) 2 weeks post-transduction (Figure 4A). This is contrary to our previous observation in 3D spheroids where successful viral transduction was limited to a small population of neurons near its outer surface (Figure 1I). To track microglia in mixed cultures with neurons, we pre-labeled iMGLs with the microglia-specific dye Isolectin IB4 (Boscia et al., 2013) (Figure 4A and Supplemental Movie 3). We found that iMGLs persisted in these mixed cultures for at least 4 weeks, with a subset of these cells adopting highly ramified morphologies shown by immunostaining with the microglial marker IBA1 (Figure 4B).

**Figure 4.**
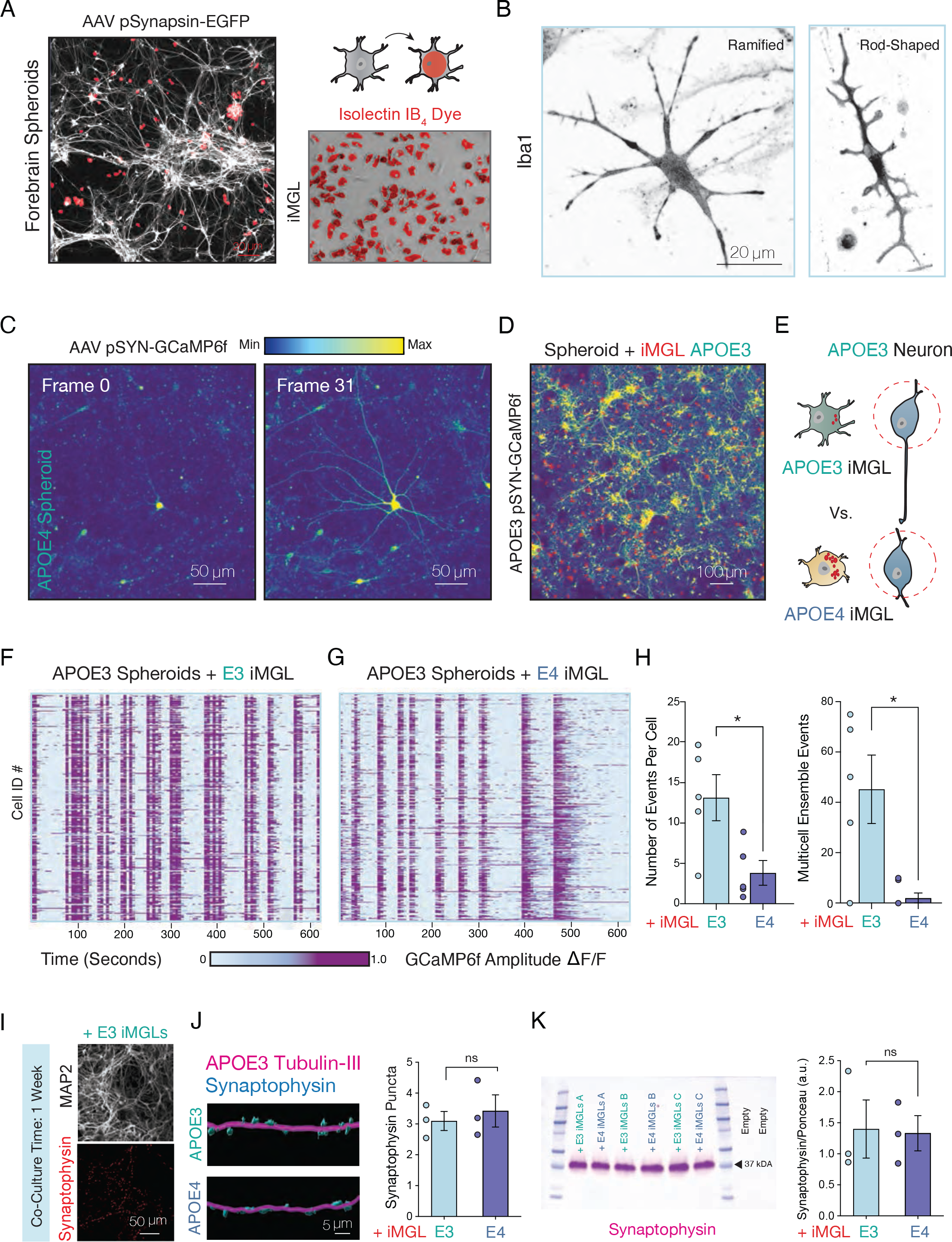
APOE4 iMGL Disrupts the Synchronized Coordination of Neuronal Ensembles. (A) APOE3 spheroids were enzymatically dissociated into a single-cell suspension and plated on laminin-treated coverslips before day 60 and cultured for an additional 4 weeks. Cultures were infected with AAV carrying EGFP under the synapsin promoter (pSYN) to label neurons. APOE3 (shown in this panel) or APOE4 iMGLs that were pre-labeled with Alexa Fluor 594 isolectin GS-IB4 conjugate for cell-tracking (red) were seeded to create a neuron-microglia co-culture model. (B) iMGLs in co-culture with spheroids adopt a highly ramified morphology shown by immunostaining with the microglia marker, IBA1. (C) Activity of a single neuron from APOE4 spheroid cultures shown over 31 frames of 3 seconds each. (D) Dissociated spheroids infected with AAV pSYN-GCaMP6f in co-culture with iMGLs labeled by isolectin in red; maximum intensity projection of the calcium indicator GCaMP6f expressed by neurons shown. (E) Diagram depicting experimental approach to measure calcium transients in neurons, while in co-culture with APOE3 or APOE4 iMGLs. (F) Spontaneous neuronal network events plotted for APOE3 neurons in co-culture for 1 week with either APOE3 or (G) APOE4 iMGLs over 10 minutes shown as raster plots. Each row corresponds to a unique neuron within the recording field-of-view, while the x-axis represents time. A threshold of 0.65 ΔF/F was used to define events of neuronal activity shown in deep purple. (H) Average number of events per cell and number of coordinated calcium transients with greater than 60% co-active cells indicative of ensemble events were quantified. Unpaired t-test; * p-value <0.05; n= 50-106 cells per group for 5-6 distinct experiments. (I) Co-culture of iMGLs with APOE3 neurons. MAP2-positive neuronal cultures exhibit synaptic puncta as evident by synaptophysin staining. (J) IMARIS 3D reconstruction of confocal z-stacks from neuron-microglia co-cultures from APOE3 neurons with either APOE3 and APOE4 iMGL immunostained for the neuronal marker Tubulin-III in magenta, and the synaptic marker Synaptophysin in blue. Quantification shown for number of synaptophysin-positive puncta per 100μm of tubulin-positive neuronal filament. Unpaired t-test; ns = not significant; n= averages from 3 biological replicates quantified from 3 field-of-view per replicate using IMARIS. (K) Western blot analysis of APOE3 spheroids in co-culture with APOE3 or APOE4 iMGL for 1 week for synaptophysin also shows no change in global synaptic content between genotypes. Unpaired t-test; ns = not significant; n= 3 separate experiments.

We next derived cortical spheroids and iMGLs in parallel from APOE3 or APOE4 isogenic iPSCs for combinatorial experiments to mix and match APOE genotypes. Dissociated spheroids from APOE3 or APOE4 were transduced with AAV pSyn-GCaMP6f for 2 weeks, and calcium dynamics visualized (Figure 4C). We observed vastly different baseline calcium dynamics between APOE3 and APOE4 neurons; while APOE3 neurons displayed highly synchronous network events, calcium transients in APOE4 neurons were asynchronous and more frequent (Supplementary Figure S3E and Supplemental Movie 4). Given the critical role of microglia in sculpting neural dynamics, we sought to determine the impact of APOE4 iMGLs to neuronal network activity. Isolectin-labeled APOE3 or APOE4 iMGLs were seeded with distinct cultures of APOE3 dissociated spheroids that had been pre-labeled with AAV pSyn-GCaMP6f prior to co-culture (Figure 4D and Figure 4E). By quantifying spontaneous calcium events of APOE3 neurons in co-culture for 1 week with either APOE3 (Figure 4F) or APOE4 iMGLs (Figure 4G), we found that APOE4 iMGLs decreased the overall number of calcium transients in APOE3 spheroid cultures (Figure 4H). Additionally, we observed that APOE4 iMGLs disrupted coordinated neuronal ensemble events in APOE3 neurons, as quantified by the number of spontaneous calcium transients with greater than 60% co-active cells (Figure 4H). We did not detect differences in the number of synapses via immunostaining (Figure 4I-J) or Western blotting (Figure 4K) for the pan-presynaptic marker, synaptophysin, after 1 week of co-culture. Since we observe changes to neuronal calcium dynamics at a point in which we do not detect robust changes to synaptic number, we postulated that a non-phagocytic mechanism may mediate the contribution of APOE4 iMGLs to impaired neuronal network dynamics.

### Imbalance in the net flux of lipids by APOE4 iMGLs

Microglia are known to secrete a vast array of immunological factors that can modulate the survival and proliferation of cells residing in neurogenic niches (Aarum et al., 2003; Kreisel et al., 2019; Ribeiro Xavier et al., 2015). To investigate if microglial secreted factors impact neuronal activity, we next decided to conduct media carry-over experiments from APOE3 or APOE4 iMGL monocultures to APOE3 spheroid cultures labeled with AAV pSyn-GCaMP6f (Figure 5A). iMGLs were incubated in unspent neuronal media for 24 hours before media was collected to ensure media carry-over would be the least disruptive to neuronal cultures, and neurons were then allowed to incubate in iMGL conditioned media for an additional 24 hours. We observed a robust decrease in neuronal calcium transients in cultures that were exposed to APOE4 iMGL conditioned media, while cultures treated with APOE3 iMGL conditioned media continued to display highly synchronized calcium transients (Figure 5B, Figure 5C and Supplemental Movie 5). Although these results were in agreement with our previous co-culture experiments, the magnitude of the neuronal activity suppression appeared to be much larger. This might be potentially due to the acute nature of the experimental design, as neurons in co-culture with microglia for several days may adapt to enriched microglial-secreted factors by modulating the expression of surface receptors. Nevertheless, we reasoned that we could take advantage of this system to dissect the mechanism by which APOE4 iMGLs differentially impact neuronal activity.

**Figure 5.**
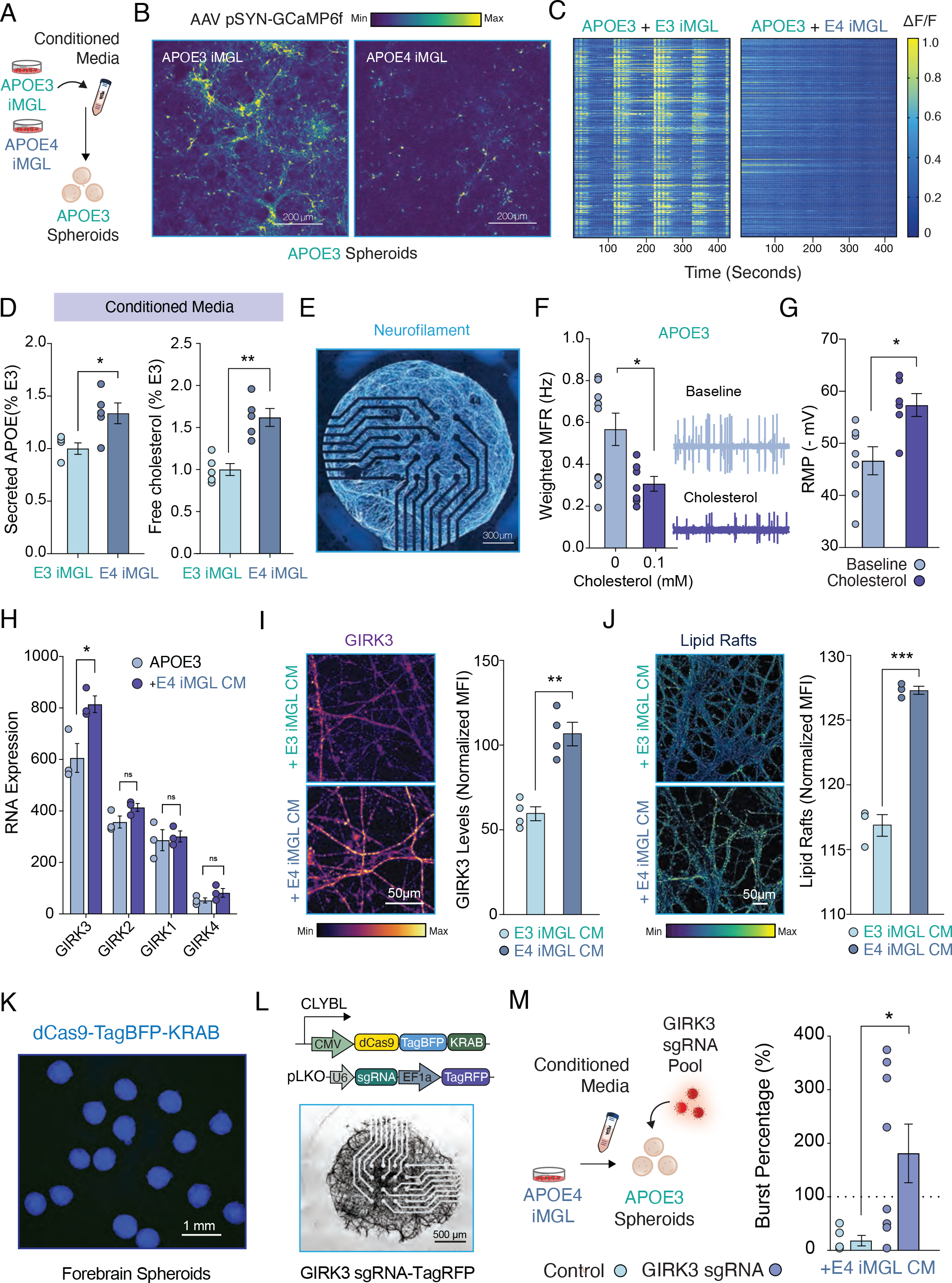
Conditioned Media from APOE4 iMGLs is Sufficient to Disrupt Neuronal Activity via Potentiation of Neuronal Lipid-Gated K+ Currents. (A) To define if neuronal calcium dynamics are altered by soluble factors secreted by iMGLs, we applied iMGL conditioned media (CM) onto APOE3 neurons expressing pSYN-GCaMP6f. (B) Dissociated spheroid cultures of 90 days were treated with iMGL conditioned neuronal media for 24 hours, and calcium dynamics analyzed over 7 minutes. (C) Heatmap for changes in GCaMP6f fluorescence intensity for APOE3 spheroids presented with APOE3 or APOE4 iMGLs CM. (D) Elisa from conditioned media of iMGLs in monoculture reveals accumulation of secreted APOE and Cholesterol. Unpaired t- test, ** p-value <0.01; * p-value <0.05. n= Averaged results of technical replicates from media sampled from 5 distinct biological replicates. (E) NGN2-induced neurons plated on a multielectrode array (MEA) immunostained for neurofilament and shown in cyan to white based on signal intensity. (F) Weighted mean firing rate (MFR) for NGN2-neurons induced from APOE3 in the presence of cholesterol shows reduced neuronal activity. MEA traces are shown for a random 120 seconds of the total 15 minutes recording session from which the data in quantified. n= 10 wells from 0mM, and 7 wells from 0.1 mM. Unpaired t-test; * p-value <0.05. (G) Patch-clamp electrophysiology of APOE3 neurons; Cholesterol treatment hyperpolarizes the resting membrane potential (RMP) of APOE3 neurons. (H) RNA-seq analysis of APOE3 spheroids at 90 days shows the lipid-gated inwardly rectifying K+ channel GIRK3 is the most abundant GIRK subunit expressed in these human-derived cultures, and its expression is elevated upon incubation with iMGL APOE4 conditioned media. n= 3 biological replicates. (I) APOE3 neurons from dissociated spheroids were incubated with APOE3 or APOE4 iMGL conditioned media for 48 hours, and analyzed for GIRK3 surface level expression by immunostaining. Representative images of fluorescence intensity are shown in a color gradient from low (purple) to high (yellow), and the mean fluorescence intensity (MFI) normalized for total arbor length in the field of view used for quantification. Unpaired t-test, ** p-value <0.01. n= averages from 3 fields of view imaged from 3 distinct biological replicates per group. (J) APOE3 neurons from dissociated spheroids were incubated with APOE3 or APOE4 iMGL conditioned media for 48 hours, and analyzed for levels of membrane lipid microdomains (also known as lipid rafts) using a cholera toxin subunit B reagent labeled with green fluorescence. Representative images of fluorescence intensity are shown in a color gradient from low (purple) to high (yellow), and the mean fluorescence intensity (MFI) normalized for total arbor length in the field of view used for quantification. Unpaired t-test, *** p-value <0.001. n= averages from 3 fields of view imaged from 3 distinct biological replicates per group. (K) iPSCs harboring a knock-in of dCAS9 fused to the repressor KRAB and tagged with blue fluorescent protein (TagBFP) induced into forebrain spheroids. (L) Neurons generated from a PiggyBac transposon-based NGN2 inducible stable line generated from CRISPRi line, seeded on MEA and transduced with a pool of 3 distinct GIRK3 sgRNAs expressing a tagged red fluorescent protein (TagRFP) reporter. (M) GIRK3 CRISPRi or APOE3 neurons infected with a non-targeting sgRNA control were seeded on MEAs and exposed to APOE4 iMGL CM for 48 hours. Dotted line represents baseline recording. Unpaired t-test, * p-value <0.05. n= Averages per well for 5-8 distinct biological replicates.

Our transcriptional profiling revealed a dramatic difference in lipid metabolism between APOE3 and APOE4 iMGLs that was associated with an activated state distinct from homeostatic surveillance. As the major transporter of cholesterol in the brain, APOE mediates the delivery of cholesterol and other lipids between neurons and glia (Ioannou et al., 2019). Thus, we decided to examine levels of APOE and cholesterol in the supernatant of APOE3 and APOE4 iMGL monocultures. We found that the supernatant of APOE4 iMGL cultures were enriched in both APOE and cholesterol in relation to APOE3 (Figure 5D). We repeated this experiment using the fluorescent cholesterol analog BODIPY-cholesterol, which also revealed an increase in cholesterol in the media of APOE4 iMGL (Supplementary Figure S4A-B). Although our transcriptional profiling is in alignment with recent suggestions that APOE4 glia exhibits reduced lipid transport (Julia TCW, 2019) (Supplementary Figure S4C), we reasoned that extracellular lipid accumulation could also be the net product of a relatively greater deficit in lipid influx. To test this idea, we exposed APOE3 or APOE4 iMGLs to low-density lipoprotein (LDL) isolated from human plasma. Along with APOE, lipoproteins like LDL make up the structural outer surface of lipid cores that are secreted from glia and transported to neurons via lipoprotein binding receptors. We observed a dramatic reduction in cellular uptake of LDL by APOE4 iMGLs in comparison to APOE3 controls (Supplementary Figure S4D-F). These results are consistent with our transcriptional profiling, in which the low-density lipoprotein receptor (LRP1) is significantly down-regulated in APOE4 iMGLs (As shown in Figure 3G). We concluded that extracellular lipid accumulation is likely a reflection of a greater impairment in lipid influx than efflux in APOE4 iMGLs already burdened by high intracellular lipid content.

To define how microglial lipid metabolism might regulate neural network dynamics, we next turned to neurons seeded onto multielectrode arrays (MEA) (Figure 5E). Seeding intact APOE3 spheroids onto MEAs yielded a robust readout of neuronal activity (Supplementary Figure S5A-B). Exposing intact APOE3 spheroids to APOE4 iMGL conditioned media for 24 hours significantly decreased the overall number of neuronal spikes and bursts relative to cells exposed to APOE3 iMGL conditioned media (Supplementary Figure S5C). Interestingly, APOE3 spheroids can partially recover after withdrawal of APOE4 iMGL conditioned media, suggesting neuronal activity is actively suppressed by soluble factors in the media (Supplementary Figure S5D).

### Neuronal accumulation of cholesterol-enriched membrane microdomains modify K+ currents

Cholesterol is essential for proper neuronal physiology, and cholesterol depletion is known to impair neurotransmission (Linetti et al., 2010; Liu et al., 2010). On the other hand, increased cholesterol efflux is well-established to enhance amyloidogenesis (Di Paolo and Kim, 2011), and decreasing cholesterol biosynthesis in astrocytes was found to alleviate plaque formation and tau burden in AD mouse models (Wang et al., 2021). Since we detect extracellular accumulation of cholesterol in APOE4 iMGLs, we wondered what the impact of exogenous cholesterol treatment would be to these neuronal cultures. Due to the extended culture times of spheroids, we opted to conduct these studies with excitatory neurons generated by the rapid induction of NGN2 expression as previously established (Zhang et al., 2013). Treatment of neurons seeded onto MEAs with water-soluble cholesterol phenocopied our observation of treatment with APOE4 iMGL conditioned media (Figure 5F). To further dissect the physiological process that renders neurons less excitable upon treatment with exogenous cholesterol, we next assessed iPSC-derived neurons by patch-clamp electrophysiology. We decided to do this work with dissociated spheroid cultures since they displayed mature neuronal calcium dynamics, perhaps due to the extended culture time in 3D before dissociation. Indeed, current-clamp and voltage-clamp recordings demonstrated physiological properties akin to mature neurons in these cultures (Supplementary Figure S3B-C). We found that addition of exogenous cholesterol significantly hyperpolarized the resting membrane potential (RMP) of cholesterol-treated neurons (Figure 5G). Moreover, we also observed a change in the I-V (Current-Voltage) curve of non-treated versus cholesterol treated neurons, indicative of greater inwardly-rectifying potassium (Kir) currents (Supplementary Figure S3D). Potentiation of Kir currents is aligned with our observation in cells recorded on MEA, as strong inwardly-rectifying potassium currents are known to hyperpolarize resting membrane potentials and decrease neuronal excitability (Hodge, 2009). In fact, overexpression of the inwardly-rectifying K^+^ channel, KIR2.1, has been used extensively in neuroscience to genetically inhibit neuronal activity (Johns et al., 1999).

Nevertheless, members of the Kir-family of channels span across 7 subfamilies (Kir1-7), and their expression and function remain uncharacterized in forebrain spheroids. We decided to isolate mRNA from APOE3 spheroids and APOE3 spheroids exposed to APOE4 iMGL conditioned media for 24 hours for bulk RNA-sequencing to define the expression of Kir channels. We detected transcripts for 14 out of the 15 annotated genes that encode Kir channel members in our forebrain spheroids, with the most abundantly expressed member being Kir3.3 (encoded by the gene KCNJ9) (Figure 5I and Supplementary Figure S3F). Kir3.3 is a G protein-gated inwardly rectifying K^+^ channel (GIRK3) that regulate neuronal excitability similarly to other Kir channels, with gain of function reducing neuronal activity and loss of function increasing neuronal activity (Luscher and Slesinger, 2010). Notably, GIRK channels are known to be lipid-gated as K+ flux is dependent on binding of the phospholipid phosphatidylinositol (4,5)P2 (PIP2) (Huang et al., 1998). In neurons, cholesterol has been shown to modulate GIRK currents, with cholesterol enrichment enhancing channel activity and thus decreasing neuronal excitability (Bukiya et al., 2019; Bukiya et al., 2017; Mathiharan et al., 2021). Based on our RNA-seq analysis, APOE4 iMGL conditioned media significantly upregulated transcript levels of GIRK3 in spheroids, while GIRK1, GIRK2 and GIRK4 remained unchanged (Figure 5I). To determine if GIRK3 upregulation was APOE genotype-dependent, we next assessed expression of GIRK3 in APOE3 spheroids exposed to either APOE3 or APOE4 iMGL conditioned media by immunostaining. Protein levels of GIRK3 (normalized for total neuronal content imaged with the pan-neuronal marker TUJ1) was significantly increased in spheroid neurons treated with APOE4 iMGL conditioned media, suggesting greater surface expression levels (Figure 5J). Interestingly, GIRK channels are known to localize to cholesterol-rich microdomains at neuronal membranes, often referred to as lipid rafts (Delling et al., 2002). It is thought that localization of receptors to lipid rafts can influence the potency of receptor-activated signaling cascades (Allen et al., 2007). Congruent with our observation of increased GIRK3 in neurons treated with APOE4 iMGL conditioned media, we detect an increased prevalence of cholesterol-rich lipid rafts in contrast to neurons treated with APOE3 iMGL conditioned media (Figure 5H).

To determine if the potentiation of GIRK3 is necessary for the suppression of neuronal activity in APOE4 iMGL conditioned media treated cultures, we next targeted GIRK3 with CRISPR-interference (CRISPRi). We derived spheroids from iPSCs edited to harbor a catalytically dead CAS9 (dCAS9) fused to the transcriptional Kruppel-associate box (KRAB) repressor within the safe harbor locus, CLYBL (Figure 5K). Gene repression in iPSC-derived neurons with this CRISPRi vector has been previously reported (Tian et al., 2021), and iPSC line edited to carry this construct was acquired from the Allen Institute Cell Collection. We determined the APOE genotype of this iPS donor to be homozygote for APOE3 in-house (data not shown). Neurons derived from these cultures were seeded onto MEAs and transduced with a lentiviral vector carrying sgRNA and a red fluorescent protein tag (TagRFP) (Figure 5L). APOE3 neurons infected with a lentiviral pool of 3 distinct sgRNAs targeting KCNJ9 (Gene that encodes GIRK3) after 3 weeks were exposed to APOE4 iMGL conditioned media. We determined by qPCR for KCNJ9 that this viral pool repressed KCNJ9 expression by 66% ± 13.9% S.E.M relative to control (vector free of targeting sgRNAs was used as control). Interestingly, we observed that GIRK3 knock-down prevented the repression of neuronal bursts by APOE4 iMGL condition media (Figure 5M). Of note, we did not observe a significant rescue in the number of spikes between GIRK3 knockdown and control (unpaired t-test, p-value = 0.1758), suggesting GIRK currents may be particularly important to regulate neuronal burst firing properties as previously demonstrated to be the case in pacemaker neurons (Li et al., 2013). Collectively, these results demonstrate that extracellular cholesterol accumulation, at least in part due to poor lipid re-uptake by APOE4 iMGLs, can suppress neuronal activity via potentiation of GIRK currents.

### Modulation of intracellular lipid content can reversibly drive purinergic signaling in microglia

To determine if APOE4-induced lipid accumulation is necessary and sufficient to drive microglia activation status away from homeostatic surveillance, we attempted to bidirectionally modulate lipid content in APOE3 or APOE4 iMGLs. We began by inducing lipid droplet accumulation in APOE3 iMGLs by exposing the cells to the mono- unsaturated fatty acid, oleic acid (OA) for 16 hours (Figure 6A). Fatty acid overload is a potent inducer of lipid droplet formation, and as such IBA1-positive cells treated with OA accumulated intracellular BODIPY-positive lipid droplets (Figure 6B-C). The rise in lipid droplet content was also linked with a decrease in the cell size of iMGLs (Figure 6D), which resembles ameboid-like morphologies adopted by activated microglia. In fact, profiling OA-treated iMGLs by qPCR for the pro-inflammatory MHC-II marker, CD74, reveals upregulation of this gene which more closely resembled basal levels observed in non-treated APOE4 iMGLs. To determine if OA-treated iMGLs exhibited deficits in purinergic signaling that phenocopied APOE4 iMGLs, we turned to calcium imaging upon ATP uncaging (Figure 6F). OA treatment was sufficient to blunt calcium transients evoked by ATP uncaging relative to untreated APOE3 iMGLs. A key mechanism mediating lipid storage into intracellular droplets is the activation of fatty-acids by the Acyl-Coa synthethase, ACSL1 (Ellis et al., 2010; Stremmel et al., 2001). ACSL1 expression has been reported to be modulated by lipogenic conditions (Li et al., 2006), and as such we observed that OA significantly induced the expression of ACSL1 (Figure 6G). This is of particular interest, since we also uncovered ACSL1 as the most enriched gene governing lipogenesis in APOE4 iMGLs through our RNA-seq analysis (shown in Figure 3I). Collectively, these results suggest that increasing lipid accumulation is sufficient to shift microglia away from homeostatic surveillance, and phenocopies key aspects of the APOE4 iMGL state.

**Figure 6.**
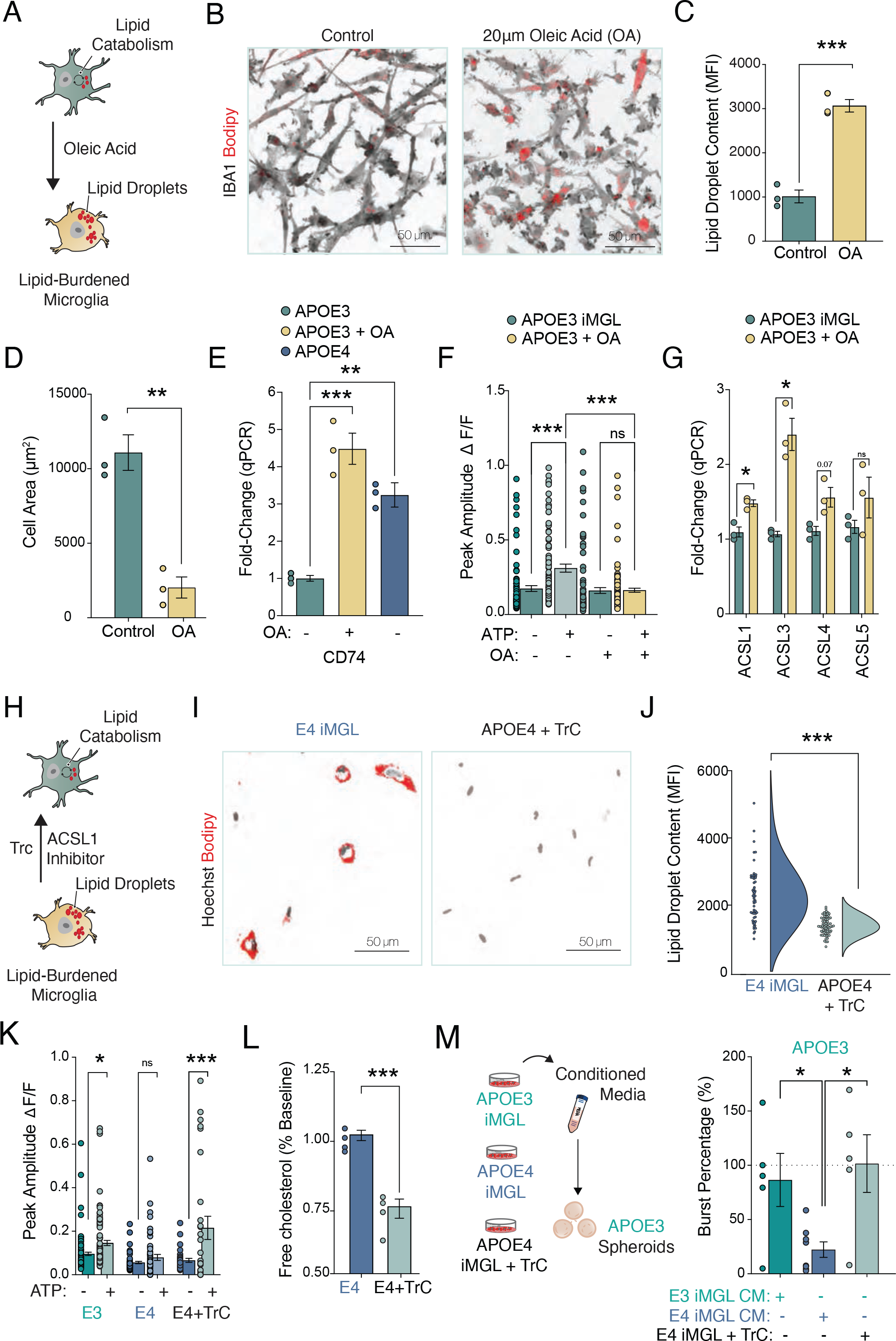
Bidirectional Manipulation of Lipid Content Can Reversibly Drive Purinergic Signaling in iMGLs. Inducing lipid accumulation in APOE3 iMGL, phenocopies APOE4 iGML’s blunted response to ATP, while depleting lipid accumulation in APOE4 iMGL can restore purinergic signaling. (A) iMGLs derived from a healthy control donor treated with 20μM of Oleic Acid (OA) for 12 hours adopt an activated morphology, shown in (B) and quantified in (C). Unpaired t-test; ** p-value <0.01; n= Averages from 3 biological replicates in each group with 20-30 cells quantified per replicate. Controls were treated with 0.05% BSA as vehicle. (D) Oleic Acid induces accumulation of lipid droplets, quantified in (E). BODIPY staining shown in red, IBA1 in gray. Unpaired t-test; *** p-value <0.001; n= Averages from 3 biological replicates in each group with 20-30 cells quantified per replicate. Data presented as grouped average per cell quantified in each group. MFI= Mean fluorescece intensity. (F) qPCR analysis for CD74, an MHC class II component associated marker of inflammation, for APOE3 iMGLs treated with vehicle or 20μM of Oleic Acid (OA). ANOVA with post-hoc Tukey test. *** p-value <0.001; ** p-value <0.01. n= averages of 3 technical replicates from 3 biological replicates. Data normalized to GAPDH expression levels and shown relative to APOE3 control. (G) OA treatment blunts iMGL response to ATP uncaging, phenocopying APOE4 iMGL. Peak amplitude of calcium transient observed with live imaging and FLUO-4 AM. ANOVA with post-hoc Tukey test. n.s. = not significant; *** p-value <0.001. n= 85-110 cells quantified per group. Experiment repeated 3 times. (H) qPCR analysis for a family of long-chain-fatty-acid-CoA ligases (ASCL1 – 5) in IMGLs shows upregulation of ASCL1 and ASCL3 in response to OA. (I) Treatment of iMGLs with 1μM of Triacsin C for 16 hours depletes lipid droplets in iMGLs. (J) Raincloud plots for lipid droplet content presented as BODIPY mean fluorescence intensity (MFI) for APOE4 iMGL treated with DMSO (vehicle) in blue or 1mM Triacsin C (APOE4 + TrC) in yellow after 16 hours of drug treatment. Unpaired t-test; *** p-value <0.001. (K) Reducing lipid droplets in APOE4 iMGLs with Triacsin C treatment is sufficient to restore response to uncaging of 1mM ATP. n= 29-52 cells per group; ANOVA with post-hoc Tukey test. n.s. = not significant; ** p-value <0.01; *** p-value <0.001. (L) Treatment of iMGLs with 1μM of Triacsin C for 16 hours, and media sampled 24 hours post drug withdrawal shows marked reduction in extracellular cholesterol via Elisa. Unpaired t-test, *** p-value <0.001; n= Averaged results of technical replicates from media sampled from 4 distinct biological replicates. (M) APOE3 neurons on MEAs exposed to conditioned media (CM) from iMGLs treated with 1μM of Triacsin C for 16 hours, and allowed to incubate for additional 24 hours post drug withdrawal. MEA recording conducted 48 hours post CM incubation onto neurons. Unpaired t-test, * p-value <0.05. n= Averages per well for 5- 8 distinct biological replicates.

Having induced lipid formation in APOE3 iMGLs, we next attempted to deplete APOE4 iMGLs of lipid droplets to test whether this would alleviate APOE4 phenotypes. We turned to the ACSL1 inhibitor Triacsin C (TrC), a pharmacological intervention that has been shown to prevent intracellular lipid accumulation in astrocytes and microglia (Marschallinger et al., 2020; Qi et al., 2021) (Figure 6H). Treatment of APOE4 iMGLs with TrC for 16 hours was sufficient to dramatically reduce BODIPY-positive lipid droplets relative to DMSO-treated control cells (Figure 6I-J). Furthermore, purinergic signaling was restored after lipid droplet depletion in APOE4 IMGLs (Figure 6K). Since TrC has also been reported to inhibit cholesterol biosynthesis and increase microglial phagocytosis (Igal et al., 1997; Marschallinger et al., 2020), we decided to test if the extracellular accumulation of cholesterol in APOE4 iMGLs could be reduced by TrC treatment. After 16 hours, DMSO-treated controls or TrC-treated APOE4 iMGLs were washed off the drug treatment and allowed to condition the media for an additional 24 hours. Indeed, we observed a significant decrease in the levels of accumulated cholesterol in the media via ELISA (Figure 6L). We reasoned that a decrease in extracellular cholesterol accumulation in APOE4 iMGLs cultures treated with TrC was likely to also relieve the suppression of neuronal activity under APOE4 iMGL conditioned media. NGN2-induced neurons from APOE3 or APOE4 iPS lines were seeded onto MEAs and allowed to mature for 3 weeks before being recorded 24 hours after exposure to conditioned media from APOE3 iMGLs, APOE4 iMGLs, or APOE4 iMGL treated with TrC (Figure 6M). While APOE4 iMGL conditioned media decreased neuronal bursts, as we previously observed, pre-treatment of APOE4 iMGLs with TrC increased neural activity to the level of neurons treated with APOE3 iMGL media. These results establish that the maintenance of lipid homeostasis in microglia sustains surveillance homeostatic states required to support proper neuronal network function. We therefore define cholesterol metabolism as a critical link between the immunometabolism of microglia and the regulation of neuronal activity.

## Discussion

### Lipid Accumulation and the Shift in Microglial Homeostatic State

Our data suggests that the inability to break down lipids is a major contributor to its intracellular accumulation in APOE4 harboring microglia. Through a process known as lipolysis, lipids are broken-down into free fatty acids to be used as energy substrates via mitochondrial oxidation (Ralhan et al., 2021). In our transcriptional profiling, we uncovered that APOE4 expression in microglia induces a switch in energy production from fatty acid oxidation to glycolysis that is accompanied by drastically reduced expression level of lipid catabolism genes. For example, APOE4 iMGLs showed strong reduction in the expression levels of Lipase A (LIPA), which is known to catalyze the hydrolysis of cholesteryl ester and triglycerides (Figure 3G). Mutations that reduce the activity of LIPA are associated with cholesteryl ester storage disease (Ries et al., 1998). In mice, LIPA deficiency leads to system metabolic and immune deficits, including the severe accumulation of intracellular lipids, increased usage of glucose as energy substrate, and inflammation (Li and Zhang, 2019). We have previously reported that APOE4 iMGLs displayed decreased motility, compromised amyloid uptake and reduced ramified morphology (Lin et al., 2018). Here we also demonstrate that APOE4 induces an activated state in microglia that in addition to lipid droplet accumulation and poor uptake of lipids and fatty acids, is marked by downregulation of canonical homeostatic markers, such as P2RY12, CSF1R and CX3CR1 and displayed impaired purinergic signaling.

Since purinergic signaling provides a window into the cellular state of microglia (i.e. P2RY12 expression is maintained by homeostatic microglia while repressed by activated microglia), we leveraged functional ATP uncaging assays to determine that the accumulation of intracellular neutral lipid stores is necessary and sufficient to shift APOE4 microglia state away from homeostatic surveillance.

### Linking Lipid Metabolism and Inflammation

Despite the innocuous nature of the stimulus, APOE4 iMGLs exhibit a pro-inflammatory response to neuronal cues. Although the substrates in spheroid conditioned media that elicit this exaggerated response remains unclear, several animal models of Alzheimer’s disease have reported a common microglia phenotype where microglia become primed by the ongoing pathology (Perry and Holmes, 2014). Primed microglia respond much stronger to a secondary inflammatory stimulus than naïve microglia, and several microglial immune checkpoints mechanisms are known to restrain microglial inflammation (Deczkowska et al., 2018). Similarly, the intracellular accumulation of lipids in APOE4 may prime iMGLs and exacerbate its response to a non-toxic stimuli. Through our transcriptional analysis, we uncovered TFs that may act as immune checkpoints to curb inflammation and that fail to be induced in APOE4 iMGLs presented with neuronal media.

Counterintuitively, this pro-inflammatory phenotype in APOE4 iMGLs exposed to neuronal conditioned media was also accompanied by the upregulation of lipogenic factors, in spite of the already lipid-burdened state of these cells. This lipogenic program included the differential expression of critical regulators of lipid droplet formation (i.e. DGAT1), fatty acid metabolism (i.e. FASN and ACSL1) and sterol biosynthesis (i.e. SREBF1). With the most enriched of these factors being ACSL1 (Figure 3I). Acyl-Coa synthetases (ACSLs) are thought to direct the metabolic fate of free fatty acids into either mitochondrial oxidation or lipid synthesis (Yan et al., 2015). In adipocytes where the function of ACSLs have been mostly studied, ACSL1 has been shown to be a target gene of peroxisome proliferator-activator receptors (PPARs) (Martin et al., 1997). PPARs direct the expression of a wide array of genes that play a major regulatory role in lipid oxidation and energy production (Kersten et al., 2000). However, observations in monocytes and macrophages have suggested that ACSL1 may play a functionally distinct role in immune cells. In mouse bone marrow-derived macrophages, ACSL1 expression is induced by proinflammatory molecules and not by PPAR agonists (Rubinow et al., 2013).

On the other hand, the induction of a lipogenic program in APOE4 iMGLs exposed to spheroid conditioned media may be more similar to our observation of increased expression of ACSL1 in APOE3 iMGLs treated with Oleic Acid. Neurons produce fatty acids in an activity-dependent manner and can efflux lipids for glial transfer (Ioannou et al., 2019). Moreover, iPSC-derived neurons are also known to secrete APOE (Wang et al., 2018). Therefore, it is unclear if in our experiments ACSL1 is activated by distinct mechanisms downstream of inflammatory signaling or fatty acid overload. Nevertheless, it has been noted that ACSL1-deficient macrophages have reduced inflammatory phenotypes (Kanter et al., 2012). In our experiments, treatment of APOE4 iMGL with the ACSL1 inhibitor, Triacsin C, depletes microglia of lipid droplets and is sufficient to restore purinergic signaling which may reflect a reversal to homeostatic state. Interestingly, we observed that prolonged exposure of APOE4 iMGLs to Triacsin C was cytotoxic and therefore unlikely to serve as a therapeutic agent. This is perhaps due to the importance of lipid droplets in buffering excess lipids associated with lipotoxicity. In fact, lipid droplets are thought to safeguard cells against various types of cellular stress (Jarc and Petan, 2019). Therefore, lipid accumulation influences many processes and must be fine-tuned rather than abrogated. Notwithstanding these challenges, our studies reveal that ACSLs may act as a critical node linking inflammation and lipid metabolism.

### Searching for a Common Lipid-Burdened Transcriptional Signature

In the mouse hippocampus, aging microglia have been shown to accumulate lipid droplets, a cellular state that has been termed lipid-droplet accumulating microglia (LDAM) (Marschallinger et al., 2020). Phenotypically, LDAM is marked by reduced phagocytosis and increased secretion of pro-inflammatory cytokines, which is similar to the cellular state we have previously characterized in APOE4 bearing iPSC-derived microglia (Lin et al., 2018). Here, we further describe how the APOE4-induced microglial state is accompanied by a critical metabolic switch that is marked by decreased oxidative phosphorylation. Contrary to our findings with human APOE4 iMGLs, mouse LDAMs have been reported to exhibit increased mitochondrial fatty acid oxidation. Through gene set activity analysis, we see poor convergence of transcriptional signatures between human APOE4 iMGL and mouse LDAM (Supplementary Figure S2I). This divergence in transcriptional profiles between our datasets could be due to specie-specific differences governing microglial lipid metabolism, as it has been previously reported that lipid metabolic dysregulation is distinct between human and mouse in APOE4 glia (Julia TCW, 2019). Nonetheless, it is likely that distinct microglia states represent lipid accumulation driven by aging as in LDAM, or as a result of deficits related to lipid flux and metabolism as in APOE4. Interestingly, although LDAM shares some overlap with transcriptional signatures of microglia associated with Amyloid-*β* plaques (disease-associated microglia or DAM), the direction in the expression of these overlapping genes do not match well (Marschallinger et al., 2020). DAM microglia are highly phagocytic, and exhibit a transcriptional signature that includes the downregulation of homeostatic genes (including CX3CRR1, P2RY12, HEXB and CST3) and upregulation of lipid metabolism genes (such as LPL, TREM2, and APOE) (Keren-Shaul et al., 2017). In our study, we report a similar downregulation of homeostatic genes as DAM, but also the downregulation of lipid metabolism genes. Interestingly, LPL knock-down in mice leads to a dysfunctional microglia state that is marked by excessive lipid droplet accumulation, defective phagocytosis, impaired lipid uptake and reduced mitochondrial fatty acid oxidation (Loving et al., 2021), well aligned with our own observations in APOE4 iMGLs. Since our analysis was performed in microglia that did not face pathology, such as pre-incubation with Amyloid-*β* or Tau, it remains unclear to what degree these transcriptional signatures would change in a more complex model with multiple cell-types and in the presence of pathology. More sophisticated *in vitro* cellular models of AD that recapitulate brain pathology will be needed to address these questions.

### Neuronal Excitability and Circuit Dysfunction

In our studies, we observed that APOE4 microglia exhibit reduced capacity to uptake lipids resulting in the net accumulation of lipids extracellularly which we found to be detrimental to neuronal activity. Recently, it has been reported that co-culture of LPS activated microglia or exposure to its conditioned media impaired the mitochondrial function of neurons (Park et al., 2020). Neuronal mitochondrial metabolism is critical for activity-dependent calcium buffering, synaptic transmission and the generation of action potentials (Harris et al., 2012). Therefore, impaired mitochondrial function induced by activated microglia is likely to dampen neuronal firing similarly to our experimental observations with APOE4 iMGLs. By profiling the soluble milieu of APOE4 iMGLs in comparison to APOE3 iMGLs, we detected elevated extracellular levels of APOE and cholesterol. Since neurons have limited capacity to buffer extracellular lipids (Ioannou et al., 2019), we examined the possibility that increased cholesterol exposure resulted in the enrichment of cholesterol-rich domains at the neuronal plasma membrane known as lipid rafts. In fact, two recent studies have linked glia-derived cholesterol abundance with neuronal lipid raft expansion (Lee et al., 2021; Wang et al., 2021). In our study, we observed that treatment of neurons with APOE4 iMGLs significantly increased levels of neuronal lipid rafts and moreover, that this rise in cholesterol abundance was also associated with increased levels of the inwardly rectifying K^+^ channel, GIRK3. Through genetic perturbation with CRISPRi, we demonstrate that the activity of GIRK3 is necessary for APOE4 microglia-mediated reduction in neuronal burst firing. Although our findings might be related to neurons exposed to LPS-activated microglia, it remains to be determined how lipids secreted by APOE4 iMGL impact neuronal mitochondrial function. Although astrocytes are the primary cellular buffers of extracellular lipids and are thought to prevent the accumulation of toxic free fatty acids that are generated from neurons in an activity-dependent manner, microglia are also critical players in the regulation of lipid transport and consumption (Ioannou et al., 2019). In the context of APOE, which is predominantly expressed by astrocytes and microglia, APOE4-disrution of lipid homeostasis across both cell types is likely a double-hit on the buffering capacity of lipids within the brain, rendering neurons more susceptible to lipotoxicity. Mechanisms that decrease excitability (such as the potentiation of GIRKs) and thus the production of fatty acids to fuel the rise in metabolism associated with increased firing rates, may be neuroprotective but at the detriment of neural computations critical for learning and memory.

### Cholesterol Homeostasis

Cellular pathways that govern the regulation of lipid and cholesterol homeostasis have emerged as a central node in the pathogenesis of AD. Although brain cholesterol is synthesized locally and independent of circulating plasma cholesterol pool due to its impermeability through the blood-brain-barrier (BBB), peripheral cholesterol has been shown to enter the brain upon BBB breakdown in aged mice (Saeed et al., 2014). High levels of plasma cholesterol in mid-life have also been associated with increased risk of developing AD, however inconsistent results regarding the ability of cholesterol-lowering statins to act as neuroprotective agents have been reported (Schultz et al., 2018). The lipidation status of APOE is reduced in APOE4-harboring cells (Kim et al., 2009), which leads to a decrease in extracellular transport of lipids and it is thought to contribute to the intracellular accumulation of cholesterol (Qi et al., 2021). High intracellular levels of cholesterol due to its poor export by APOE4 may lead to incorporation of cholesterol at the mitochondrial membrane. Increased levels of cholesterol at the mitochondrial membrane is associated with decreased oxidative phosphorylation capacity and has been previously reported in mouse models of AD (Elustondo et al., 2017). Whether deficits in mitochondrial metabolic function is the cause or the effect of lipid accumulation in APOE4 microglia remains unresolved. Similarly to our findings in microglia, we have previously reported that in astrocytes derived from APOE4 iPSCs cholesterol accumulates extracellularly (Lin et al., 2018). Although the mechanism by which cholesterol and other lipids may accumulate extracellularly in APOE4 glia is not clear, astrocytes activated with pro-inflammatory stimuli have recently been reported to secrete saturated lipids contained within APOE and APOJ lipoproteins that are toxic to neurons (Guttenplan et al., 2021). Evidence exists that microglia also secrete higher levels of APOE under pro-inflammatory conditions (Lanfranco et al., 2021). Since the metabolic profile of microglia is associated with its pro-inflammatory state, therapies aimed at reprogramming microglial metabolism may prove to be imperative in curbing inflammation and halting neurodegeneration in AD.

## Acknowledgments

We thank Y.T. Lin and T. Ko for experimental assistance with iPSCs, Y. Zhou, H. Cam, M. Mazzanti and T. Garvey for administrative support. In addition to P. Narayan, L. Akay and all Tsai laboratory members for helpful discussions. Our work in the Tsai lab is only possible through the generous support of the The Belfer Neurodegeneration Consortium, Ludwig Family Foundation, JPB Foundation, Joseph P. DiSabato and Nancy E. Sakamoto, Donald A. and Glenda G. Mattes, Lester A. Gimpelson, The Halis Family Foundation, The Dolby Family, David Emmes, and Alan and Susan Patricof. This work was supported by NIH grants R01-AG058002 and RF1-AG062377 to L.-H.T. M.B.V. is supported by the Howard Hughes Medical Institute Hanna H. Gray Postdoctoral Fellowship.

## Author Contributions

M.B.V and L.-H.T. conceived the study. M.B.V., N.L., X.L., G.S. performed experiments and analyzed results. M.M. analyzed calcium imaging data, A.H.E, G.W. and M.B. provided intellectual input on experimental design and data interpretation. O.C., and H.-L.W. generated iMGLs. L.L. and N.L. performed electrophysiology analysis. H.M. performed RNA-seq analysis. D.V.M. computed gene set activity scores to the LDAM signature. L.B. performed ELISA and Western blot analysis. M.B.V and N.L. wrote the methods. M.B.V and L.-H.T. wrote and revised the manuscript.

## Lead Contact

Further information and requests for resources and reagents should be directed to and will be fulfilled by the lead contact, Li-Huei Tsai

## Materials Availability

### Data and Code Availability

All data reported in this paper will be shared by the lead contact upon request. RNA-seq data pending submission to GEO, and will be released before publication. This paper does not report original code.

## Methods

### Cell lines and differentiation from iPSCs

All human iPSCs were maintained in feeder-free conditions in mTeSR1 medium (STEMCELL Technologies) on Matrigel-coated plates (Corning; hESC-Qualified Matrix) iPSCs were passaged at 60–80% confluence using ReLeSR (STEMCELL Technologies) and reseeded 1:6 onto Matrigel-coated plates. APOE isogenic lines derived from a 75 year old female (AGO9173) with APOE3/3 genotype edited to harbor APOE4/4. A second distinct APOE isogenic line was derived from a 70 year old female sporadic AD patient (AG10788) with APOE4/4 genetoype (sADE4/4) and CRISPR-edited to APOE3/3 (sADE3/3). The iPSC lines were generated by the Picower Institute for Learning and Memory iPSC Facility as first described (Lin et al., 2018). CRISPRi iPSCs were acquired via the Allen Institute for Cell Science https://www.allencell.org, and maintained similarly as described above. APOE genotype for the CRISPRi line was determine by amplifying the APOE locus with PCR primers Forward: 5’- ATGGACGAGACCATGAAGG -3’ Reverse: 5’-CTGCCCATCTCCTCCATCC -3’ followed by Sanger sequencing with primer Forward: 5’- GCACGGCTGTCCAAGGAG -3’ and Reverse: 5’-CAGCTCCTCGGTGCTCTG -3’.

### Spheroid Induction Protocol

Dorsal forebrain spheroids were generated using a previously established protocol (Sloan et al., 2018), with an adapted iPS seeding strategy (Marton et al., 2019). Briefly, confluent iPSCs were dissociated into a single-cell suspension after incubation in ReLeSR (STEMCELL Technologies) for 2 minutes at room temperature followed by a dry incubation at 37C for 5 more minutes. iPSC colonies were then scraped in mTeSR1 medium (STEMCELL Technologies) and dissociated into a single-cell suspension by mechanical pipetting. Cell suspension was centrifuged at 300g for 5 minutes, resuspended in 1ml of mTeSR1 medium supplemented with ROCK inhibitor (Rockout, BioVision) and counted with an automated cell counter (Countess, Invitrogen). 3 x 10^6^ cells were then plated onto microwells (AggreWell 800, STEMCELL technologies) for embryoid body induction. After 48 hours, embryoid bodies were moved onto non-tissue culture treated petri dishes (Falcon, Corning) neural induction following forebrain spheroid differentiation protocol.

### Spheroid Dissociation and 2D plating

After growing in suspension for at least 60 days, spheroids were dissociated into a single cell suspension for plating onto coverslips to generate 2D cultures. Adapting a previously described protocol, spheroids were incubated in Accutase (StemPro, Life Technologies) for 30 minutes at 37C. Following Accutase aspiration, spheroids were mechanically dissociated by pipetting in Hank’s Balanced Salt Solution containing 10% FBS (HBSS, Thermo Scientific). Cell suspension was centrifuged at 300 x g, washed in warm Neurobasal media (Gibco) supplemented with B-27 (Gibco) and N-2 (Gibco) (Neuronal Media), passed through a 70μM strainer (VWR International), and plated in 24-well Poly-D-Lysine (Sigma-Aldrich) coated No. 0 glass coverslips in 6-well MatTek plates (MatTek) at a ratio of 1 spheroid per 3 wells. Cells were allowed to recover for 1 month prior to experiments in neuronal media, half-feeding every 3-4 days.

### Microglia Induction Protocol

Embryoid bodies (EBs) were generated using the same protocol as described for spheroid differentiation and seeded onto Matrigel-coated 6-well tissue culture plates at a density of 15-30 EBs per wells. EBs were first differentiated into hematopoietic progenitor cells (HPCs) using the STEMdiff Hematopoietic Kit (STEMCELL Technologies). Following a previously established protocol (McQuade et al., 2018), non-adherent HPCs were collected, centrifuged at 300 x g, and resuspended in 1 mL of microglia differentiation media (MDM) containing a mixed composition of DMEM/F12 and Neurobasal (half/half) (Gibco) supplemented with IL-34 and m-CSF (Peprotech) (Mcquade et al., 2018). Cells were plated in 6-well tissue culture plates at 200,000 cells per well and maintained in MDM for at least two weeks prior to use in experiments.

### Microglia-Neuron Co-Cultures

Dissociated neuronal cultures were switched from Neuronal Media to BrainPhys Neuronal Medium (STEMCELL Technologies) 1 month after dissociation and prior to iMGL seeding. Neurons were infected with 12μl of AAV9 hSYN-EGFP (Addgene #50465-AAV9) at titer *≥*7x10^12^ vg/mL or 12μl AAV1 SYN-GCaMP6f-WPRE-SV40 (Addgene #100837-AAV1) at titer *≥*1x10^13^ vg/mL per 24 wells. Brainphys media was reduced to 300μl overnight during transduction. The following day, fresh media was added to reach a final culture volume of 500 μL Upon harvesting and adding iMGLs, in suspension, to neuronal cultures, Brainphys was supplemented with m-CSF. Co-cultures were half-fed every 3-4 days and were ready to be used for experiments after a minimum of 1 week.

### Immunofluorescence

Spheroids, dissociated neuronal cultures, and iMGLs were fixed with 4% paraformaldehyde (Electron Microscopy Sciences) at room temperature for 20 minutes followed by three washes in Dulbecco’s PBS (Gibco). Spheroids were incubated in 30% sucrose overnight and imbedded in Tissue-Tek OTC (Sakura) for cryosectioning at 40μm slices on a cryostat (Leica CM3050s). Fixed cells or slide-mounted spheroid slices were incubated with gentle agitation for 1 hour in blocking buffer (5% BSA, 1% NGS, 0.3% Triton-X in DPBS) at room temperature. Primary antibodies were incubated in blocking buffer overnight at 4C. Secondary antibodies conjugated to Alexa-488, -555, -594, or 647 were applied at 1:1,000 for 1 hour at room temperature. Cells were incubated in Hoechst 33342 (Thermo Scientific) diluted 1:10,000 in DPBS for 5 minutes prior to mounting and imaging. Detailed list of primary antibodies used in this study can be found in the table below.

| **ANTIBODY** | **Dilution** | **Catalog Number** | **Manufacturer** |
| --- | --- | --- | --- |
| Guinea Pig Anti-IBA1 | 1:500 | #234004 | Synaptic Systems |
| Rabbit Anti-P2RY12 | 1:100 | #HPA014518 | Milipore Sigma |
| Rabbit Anti-P2X1 | 1:100 | #PA5-77662 | Thermo Scientific |
| Rabbit Anti-THIK-1 | 1:100 | #ab237624 | Abcam |
| Mouse Anti-KCNE3 | 1:100 | #sc-393841 | Santa Cruz Biotechnology |
| Rabbit Anti-CACNA2D4 | 1:50 | #A65974-020 | EpiGentek |
| Mouse Anti-GLUR7 | 1:200 | #MA5-31743 | Thermo Scientific |
| Chicken Anti-MAP2 | 1:10,000 | #PCK-554P | BioLegend |
| Mouse Anti-Tubulin β-III | 1:2,000 | #MMS-435 | BioLegend |
| Mouse Anti-SYNAPTOPHYSIN | 1:200 | #S5768 | Milipore Sigma |
| Rabbit Anti-KCNJ9 (GIRK3) | 1:100 | #HPA070478 | Atlas Antibodies |
| Mouse Anti-NEUROFILAMENT | 1:1,000 | #SMI-311R | BioLegend |

Staining with non-antibody probes were performed following manufacturer’s guidelines. Below is a detailed list of their use in this study.

| Probe | Use in the Study | Catalog Number | Manufacturer |
| --- | --- | --- | --- |
| ActinGreen ReadyProbes | To visualize cell morphology | #R37110 | Invitrogen |
| FLUO4-AM | Microglia calcium imaging | #F14201 | Invitrogen |
| BODIPY 493/503 | Lipid droplet staining | #25892 | Cayman Chemical |
| Isolectin GS-IB4 Alexa Fluor 594 | Label microglia prior to co-culture | #I21413 | Invitrogen |
| Vybrant Alexa Fluor 594 Lipid Raft Labeling Kit | Lipid raft staining | #V34405 | Invitrogen |

Microscopy was performed using a Zeiss LSM880 confocal system and fluorescent Z-stack images were quantified using IMARIS (Oxford Instruments).

### Calcium Imaging

Live-imagining was performed with Zeiss LSM900 equipped with a heated chamber kept at 37°C with humidity and CO2 control. For Fluo-4 AM labeled iMGLs, images were acquired at 488nm and compound uncaging done at 405nm for 30 seconds post baseline acquisition followed immediately by post-stimulation image acquisition. For uncaging experiments, cells were pre-incubated in 1mM Caged ATP (DMNPE-Caged ATP; Fisher Scientific #1049), 1mM Caged NMDA (MNI-Caged NMDA; Tocris #2224) or 1mM Caged Glutamate (MNI-Caged L-glutamate; Tocris #1490). Images were first stabilized to account for drift in the x-y direction, we used the ImageJ plugin “Linear Stack Alignment with SIFT”. Calcium traces from motion-corrected time series were manually segmented on ImageJ into individual cells based on threshold intensity, variance, and upper and lower limits for cell size. Image segmentation results were separately inspected for quality control. Fluorescence signal time series (ΔF/F: change in fluorescence divided by baseline fluorescence) were calculated for each individual segment whereby the baseline fluorescence for each cell was determined as the minimum fluorescence signal in baseline recording epoch. For GCaMP-tagged neurons, the onset of a calcium transient was identified as occurring when ΔF/F exceeded two standard deviations above the baseline fluorescence, and the termination of a calcium transient was identified as occurring when ΔF/F fell below 0.5 standard deviation above the baseline fluorescence. To test for changes in functional connectivity between cells in the presence of APOE3 versus APOE4 microglia, we quantified multicellular ensembles. A multicellular ensemble event was identified as occurring when the number of simultaneously active cells exceeded 60% of all cells. For neuronal calcium dynamics, data was generated from distinct cultures recorded in separate experiments and plotted as averages per group, while for iMGLs data was plotted from individual cells in one experiment, although the experiments were repeated at least three times. Heatmaps were generated using GraphPad Prism (GraphPad Software).

### Electrophysiology

Whole-cell patch-clamp recordings of neurons were performed at 6 to 8 weeks after spheroid dissociation and 2D plating, or for iMGLs after 2 to 4 weeks of iMGL differentiation. Intracellular recordings were performed at room temperature using an Axon CV-7B headstage, Multiclamp 700B amplifier, and Digidata 1440A digitizer (Molecular Devices). Electrode pipettes were pulled from borosilicate glass (World Precision Instruments) on a Model P-97 Flaming/Brown micropipette puller (Sutter Instrument) and typically ranged between 4–7 MΩ resistance. Intrinsic neuronal properties were studied using the following solutions (in mM): Extracellular: 125 NaCl, 2.5 KCl, 1.2 NaH2PO42H2O, 1.2 MgCl26H2O, 2.4 CaCl22H2O, 26 NaHCO3, 11 glucose (pH 7.4). Intracellular: 135 K-gluconate, 5 KCl, 2 MgCl26H2O, 10 HEPES, 2 Mg-ATP, 0.2 Na2GTP (pH 7.2). Membrane potentials were typically kept between - 50 mV to -70 mV depending on whether neurons or iMGLs were being recorded. In voltage-clamp mode, currents were recorded with voltage steps ranging from -160 mV to +80 mV. In current-clamp mode, action potentials were elicited by injection of step currents from -50 pA to +50 pA. For experiments aimed at determining the impact of cholesterol on neuronal properties, 1 mM cholesterol (Cholesterol Water-Soluble, Sigma-Aldrich #C4951) was supplemented to the external solution. ATP-evoked currents were recorded by local application of 100μM (Sigma-Aldrich; #A9187). Data were first collected and analyzed using pCLAMP 11 software (Molecular Devices). Further analysis was done in GraphPad Prism (GraphPad Software).

### Spheroid Electrical Stimulation

Culture pacing system (C-Pace EM, IONOPTIX) equipped with 6-well carbon electrode dishe (C-Dish, IONOPTIX) was used to deliver electrostimulation to spheroids at 12V with a biphasic pulse train frequency of 40Hz. Paddle carbon electrodes were scrubbed clean with ethanol following manufacturer’s recommended procedure and allowed to fully dry before use. To prevent effect of media hydrolysis, washes and full media switches immediately followed stimulation.

### MEA

Dissociated spheroids or NGN2-induced neurons were plated as a 10μl droplet in Poly-D-Lysine (Sigma-Aldrich) coated wells of a CytoView MEA 48-well plate (Axion BioSystems; M768-tMEA-48B). Typically 50,000-75,000 cells were plated per well that each contained 16 low-impedance PEDOT electrodes 50μm in diameter and arranged at a pitch of 350μm. Intact spheroids were plated and covered in a Matrigel droplet (Corning; hESC-Qualified Matrix) to anchor the spheroid. After 15-30 minutes in 37°C, droplets were flooded with warm Brainphys media (STEMCELL Technologies) and allowed to recover for at least 4 weeks before recording sessions. For conditioned media carry-over experiments, a recording session preceded the media treatment, denoted as baseline recording. iMGL media was added to compose half of the final volume of the well and allowed to incubate for 24 hours before a second recording was performed. All extracellular recordings were performed using the Axion Maestro Pro MEA system (Axion Biosystems). Spontaneous neural activity was recorded for 30 minutes at a sampling rate of 12.5 kHz and an adaptive threshold set at 5.5 times the standard deviation of baseline noise was used for spike detection. Bursts were detected at each electrode using an inter-spike interval (ISI) threshold set to at least 5 spikes with a maximum ISI of 100 ms. Electrodes were defined as active if neuronal firing occurred at a minimal rate of 5 spikes/min. For MEA data analysis, only wells containing a minimum of 3 active electrodes were included. Neuronal firing metrics were exported as the averages from each well from Axion Biosystems’ Neural Metrics Tool and plotted with Prism Graphpad (GraphPad Software).

### Drug Treatment

To block voltage-gated sodium channels, 1μM Tetrodotoxin (Trocris, #1078) was applied to media before electrostimulation of spheroids. 20μM Oleic Acid (Sigma-Aldrich; #03008) was applied overnight to iMGLs to induce lipid accumulation. Control cells were treated with 0.1% BSA as vehicle. 1μM Triacsin C (Cayman Chemical; #10007448) was applied to iMGLs overnight to deplete lipid accumulation. Control cells were treated with DMSO as vehicle.

### Western blot

Spheroids that had been transplanted with APOE3 or APOE4 iMGLs for 10 days were washed once with cold 1x PBS and homogenized in RIPA lysis buffer (Sigma-Aldrich, #R0278) containing Halt protease/phosphatase inhibitor cocktail and EDTA (ThermoFisher, #78440). Supernatants were collected after centrifugation at 14,500 RPM for 15 min at 4 °C and stored at -80 °C for later use. Total protein levels were quantified using the Pierce BCA Protein Assay Kit (Thermo Scientific), and 5µg of protein were loaded from each sample per lane onto precast 4-20% polyacrylamide gels (Bio-Rad, #4561094). Denatured/reduced samples were run at 150V for 75 minutes, after which proteins were transferred from the gel to 0.2μm nitrocellulose membranes (Bio-Rad, #1704159) using the Trans Blot Turbo Transfer System (Bio-Rad) set to the mixed molecular weight program. Membranes were stained with Ponceau S (CST, #59803) and subsequently blocked with 5% non-fat milk in 1x TBST (10 mM Tris-HCl pH 8.0, 150 mM NaCl, 0.05% Tween-20) for 1 hour before incubating with primary antibody. Membranes were incubated with rabbit anti-synaptophysin (CST #36406, 1:1000) overnight at 4°C, and secondary antibody was later incubated at room temperature for 2 hours. Wash buffer was 1x TBST. Proteins were detected by WesternBright Quantum HRP substrate (Advansta, #K-12042) and visualized using the ChemiDoc MP Imaging System (Bio-Rad).

Western blot densitometry was conducted using ImageJ. Synaptophysin levels were normalized to Ponceau S.

### ELISA

Cholesterol levels from iMGLs in monoculture were measured using the Cholesterol Assay Kit (Abcam #ab65390) following manufacturer’s instructions for fluorometric detection. Cells were grown in 6-well plates, and samples were either assayed immediately or frozen at -80°C Celsius and thawed once for cholesterol measurements. To obtain total cholesterol levels, cholesterol esterase was added to samples. For free cholesterol measurements, samples were used directly without the addition of enzyme. Measurements were made using an EnSpire plate reader (Perkin Elmer). APOE levels were similarly processed and quantified from iMGL conditioned media using Apolipoprotein E Human ELISA kit (Invitrogen; #EHAPOE).

### Lipid Cellular Assays

The fluorescently tagged cholesterol analog, BODIPY-Cholesterol (Cholesterol with BODIPY at carbon-24 of the side chain) (Cayman Chemical; #24618) was used to assay the extracellular accumulation of cholesterol in monocultures of APOE3 or APOE4 iMGLs. Cells were incubated with BODIPY-Cholesterol for 48 hours to saturate cellular uptake, washed 3 times and further incubated for an additional 24 hours before media was collected, centrifuged at 300g for 5 minutes and assayed for fluorescence at 488nm with EnSpire plate reader (Perkin Elmer). Low Density Lipoprotein (LDL) from human plasma complexed to pHrodo red (pHrodo-LDL) (Invitrogen; #L34356) was used to determine LDL uptake in APOE3 and APOE4 iMGLs. Monocultures were treated with 5 μg/ml of pHrodo-LDL and incubated for 1 hour before live cells were imaged with an EVOS cell imaging system (Thermo Scientific). Images were processed in IMARIS (Oxford Instruments) to reconstruct cellular boundaries and quantify intracellular content as mean fluorescence intensity.

### RNA Analysis of iMGLs and Spheroids

RNA extraction from biological replicates (n=3) for the isogenic pair (APOE3 and APOE4) of iMGLs exposed to spheroid conditioned media or unspent neuronal media was achieved with RNeasy Plus Mini Kit (Qiagen). RNA integrity number (RIN) scores were determined to be above 9 before library preparation. MIT BioMicro Center prepared libraries using the NEBNext Ultra II RNA Library Prep Kit (New England Biolabs) and performed 75 bases single-end run NextSeq 500 Illumina sequencing. FASTQ reads were aligned using STAR (v.2.6.1a) to GRCh37 reference genome (GENCODE 19) (Dobin et al., 2013). Transcripts were quantified using HTSeq, data was normalized utilizing RUV-seq and differential gene expression analysis was performed through DESeq2 as previously described (Meharena et al., 2022). Significant differentially expressed genes (DEGs) were called with an FDR < 0.05 with unrestricted log2 fold- change cut-offs. Gene ontology analysis was performed using http://bioinformatics.sdstate.edu/go/ and EnrichR Appyter https://maayanlab.cloud/Enrichr/. Gene set activity scores related to the LDAM signatures (Supplementary Table T2-1 (Marschallinger et al., 2020), padj < 0.05) were computed on the iPSC RNA-sequencing normalized counts matrix, as previously implemented in the R package GSVA (Hanzelmann et al., 2013). Briefly, GSVA first estimates gene-wise (non-parametric) Gaussian cumulative density functions based on normalized sample expression values. A KS-like random walk statistic is computed for every gene set and the enrichment score is calculated as the difference between the largest positive and negative random walk deviation from zero, which ensures that the scores follow the standard normal distribution and meet the assumptions for linear modeling. The following parameters were used to evaluate the GSVA function: mx.diff=TRUE, kcdf=c(“Gaussian”), min.sz=5. RNA extraction for qPCR analysis was performed similarly, and reverse transcription performed with RNA to cDNA EcoDry Premix (Takara) according to manufacturer’s instruction. Gene expression was analyzed with Real-Time PCR (Bio-Rad, CFX96) and SsoFast EvaGreen Supermix (Bio-Rad). Expression data was normalized to housekeeping gene GAPDH using the 2^-ΔΔCT^ relative quantification method. A list of qPCR primers used in this study can be found below.

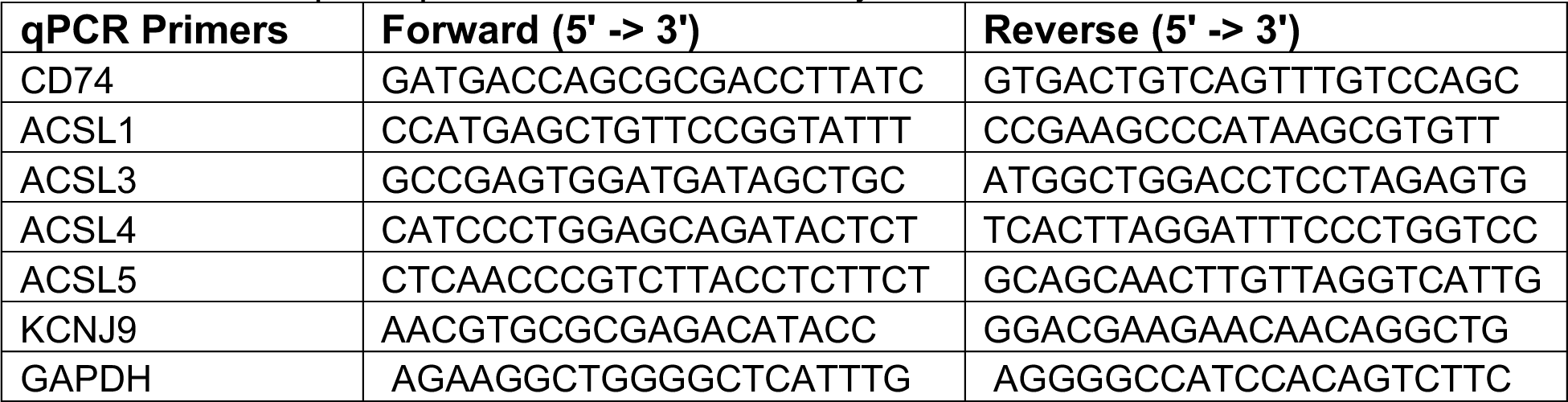

### Plasmids, Cloning and Lentivirus Production

To target KCNJ9 (GIRK3) in CRISPRi iPS-derived neurons, three distinct gene target sequences were picked from the sgRNA/gene published library for CRISPRi (Horlbeck, eLife). Protospacer sequences (note that 5’G is added independent if it exists in the genome): sgRNA 1: 5’-GCCCCCACGGGCCCCCCGAA-3’, sgRNA 2: 5’-GCACGGGCCCCCCGAAGGGT-3’, sgRNA 3: 5’ -GTGTAGCGGCAGCTCTGACT-3’. 5ug of the lentiviral vector pLKO5-sgRNA-EFS-tRFP (Addgene #57823) was linearized with BsmBI (New England Biolabs) and oligos annealed with adapters to the sense 5’ - CACC and antisense 5’-AAAC were ligated using Quick Ligase Kit (New England Biolabs). Ligated product was transformed into stabl3 competent E. coli (New England Biolabs) and transformed screened for insert via sanger sequencing with U6 forward universal primer (GeneWiz Azenta) and aligned to plasmid (SnaGene software). Clones were then transfected into HEK293Ts together with lentiviral packaging and envelope vectors to generate lentivirus following the previously published protocol (Victor et al., 2014). Lentiviral supernatant was collected 48 hours after transfection and centrifuged for 2 hours by ultracentrifugation at 25,000 RPM at 4°C (Beckman Coulter). Pellets were resuspended in DPBS and frozen in -80°C until used. All three sgRNA viruses were mixed equally to deliver a pool totaling 12ul per well of a 48-well MEA plate. Virus was top loaded and allowed to incubate overnight in neurons that had been seeded within the past 48 hours. Within 5-7 days tRFP (turbo Red Fluorescent Protein) expression was visible, and by 3 weeks the vast majority of cells in the culture expressed high levels of RFP.

### Statistics

Statistical analyses were performed in GraphPad Prism using a two-tailed Student’s t-test or a one-way ANOVA followed by a post hoc Tukey’s test with *P < 0.05 considered significant. Multiple comparisons were corrected with Dunnett method as described in the figure legends. Studies were performed blindly and automated whenever possible with the aid of IMARIS or ImageJ cell, and multiple investigators confirmed quantification results. Data in graphs are expressed as mean and error bars represent standard error of the mean (SEM). Outliers were detected and excluded with Grubbs’ test for alpha levels of 0.05. In the entirety of this study, only MEA data exhibited high variability and, therefore, had data points excluded based on this criterion. All experiments reporting values of single cells instead of population averages were performed at least three times.

## Victor_Supplemental Figure Legends

**Supplemental Figure 1.**
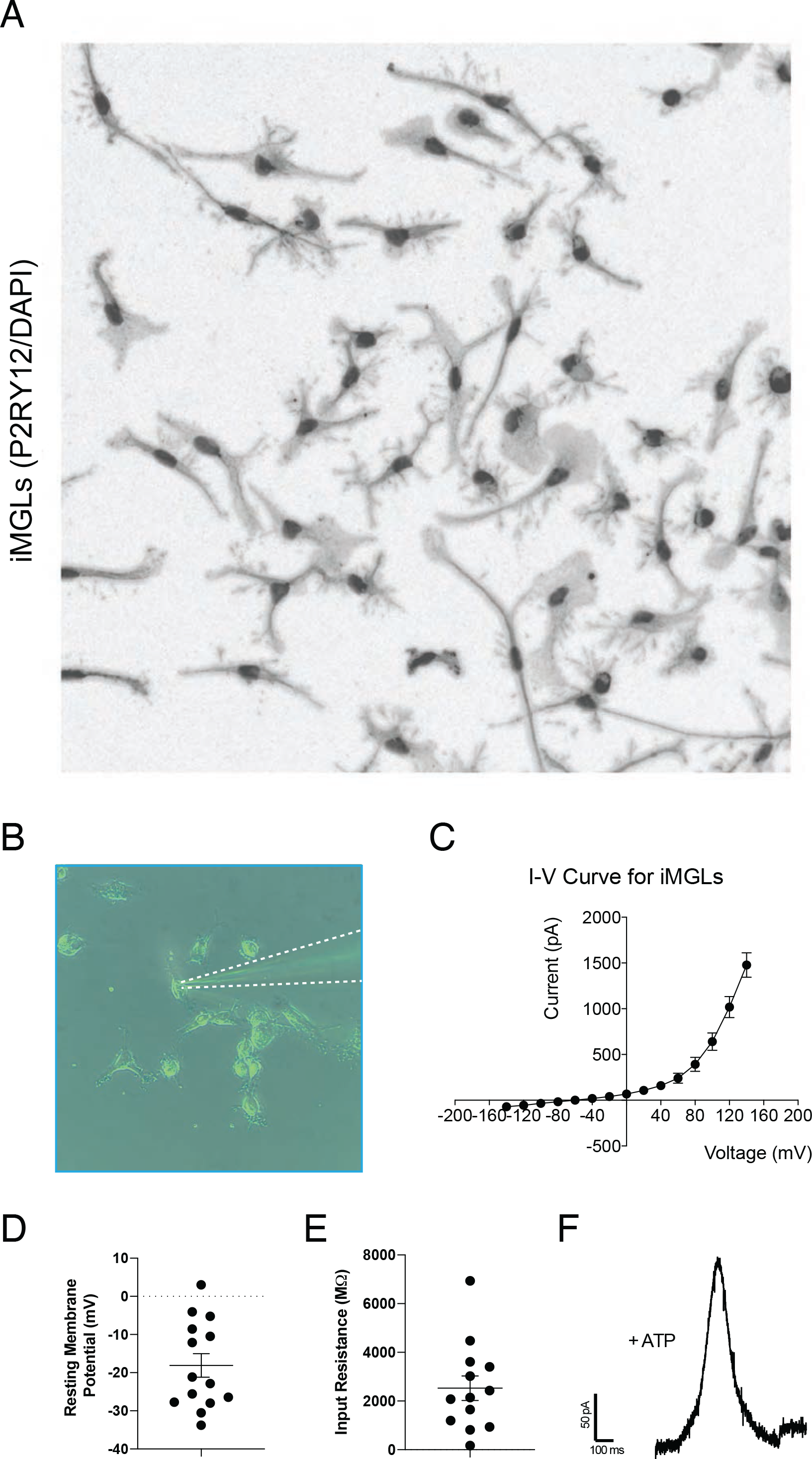
Passive Membrane Properties of iMGLs. iMGLs derived from healthy non-demented human subject were characterized with patch-clamp electrophysiology. (A) iMGLs in monoculture stained with P2RY12 and DAPI, shown as an inverted monochromatic image. Image shown as larger inset to ease morphology view from main figure 1B. (B) Image of whole cell patch-clamp recording of iMGLs in monoculture. Outline of glass electrode (5-7 MΩ pipette resistance) is shown in dashed white line. (C) Current-Voltage relationship shown for 15 cells from injection steps of 20mV ranging from -140mV to +140mV. Membrane depolarization induces large outward currents up to 1.5nA, while membrane hyperpolarization only induces small inward currents maxing at -71.13pA. (D) Resting membrane potential for recorded cells ranged from 3mV to -33mV, averaging -18mV. n=14 cells. (E) Input resistance for recorded cells ranged from 170 to 6,934 MΩ, averaging 2,527MΩ. n=13 cells.

**Supplemental Figure 2.**
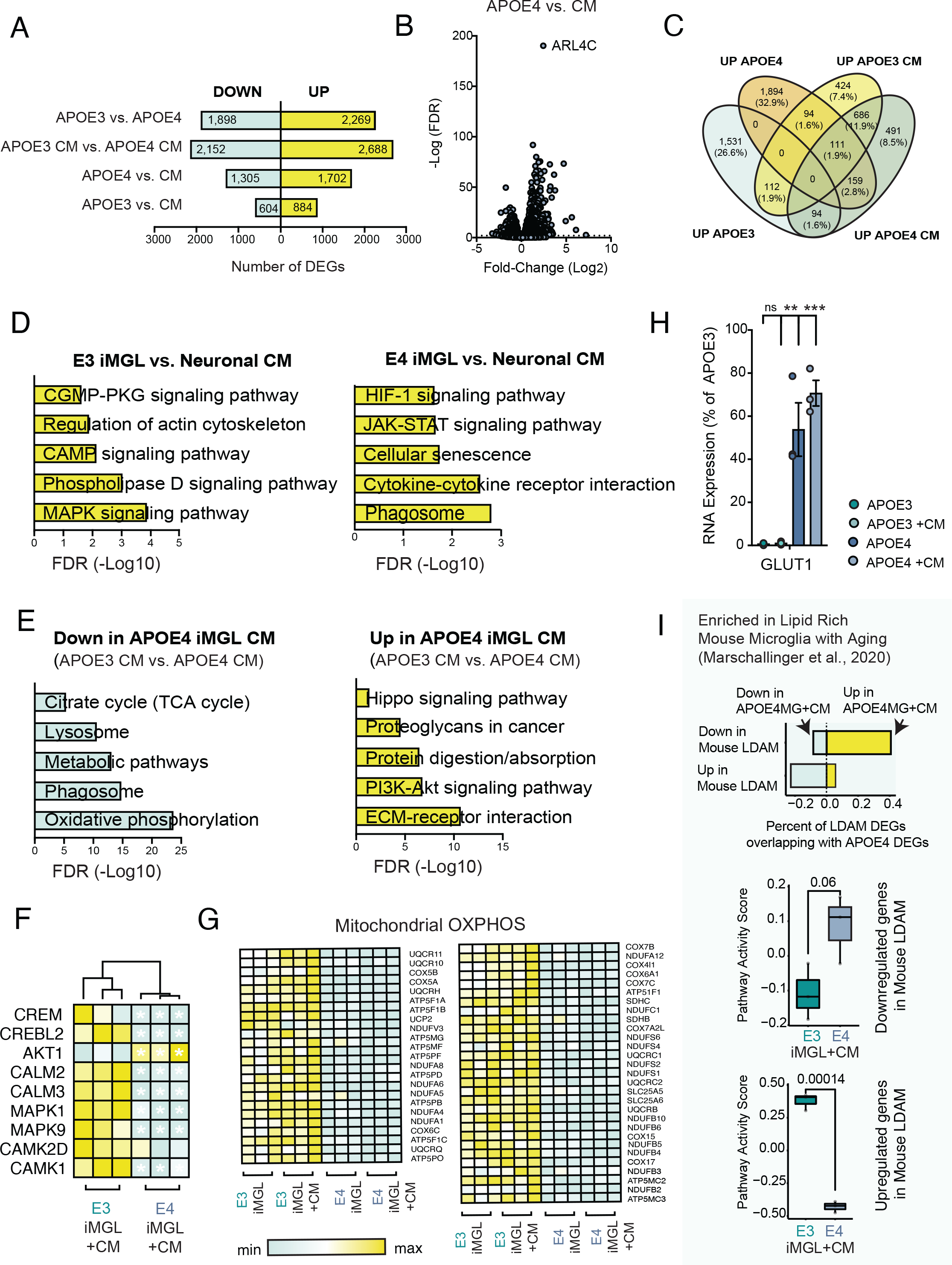
RNA-seq from iMGLs in Monoculture at Baseline or After Exposure to Neuronal Conditioned Media. (A) Histogram depicting the number of significantly differentially expressed genes (DEGs) (FDR>0.05, unrestricted cut-off for fold-change) for multiple comparisons. For each comparison (e.g. APOE3 vs APOE4), directionality of the number of DEGs are shown for the latter term (i.e. UP is highly expressed in APOE4). (B) Volcano plot for DEGs evoked by neuronal conditioned media in APOE4 iMGLs. (C) Venn diagram of up-regulated DEGs. (D-E) Curated gene ontology analysis for KEGG pathways is shown as histogram based on the FDR, the top 30 pathways ranked by p-value can be found in supplemental tables 1-4. (F) Calcium-evoked transcripts elicited by CM. (G) DEGs for mitochondrial oxidative-phosphorylation (OXPHOS) shown as a heatmap. (H) GLUT1 levels normalized to APOE3. ANOVA with post-hoc Tukey test. n.s. = not significant; ** p-value <0.01; *** p-value <0.001. (I) Overlap of down- and upregulated DEGs in APOE3 CM vs APOE4 CM microglia (FDR<0.05) and down- and upregulated DEGs in mouse lipid-low vs lipid-high microglia (LDAM) (FDR<0.05). Gene set activity scores for the Mouse LDAM downregulated and upregulated gene signature (logFC<0 and FDR<0.05) in APOE3 CM and APOE4 CM cells.

**Supplemental Figure 3.**
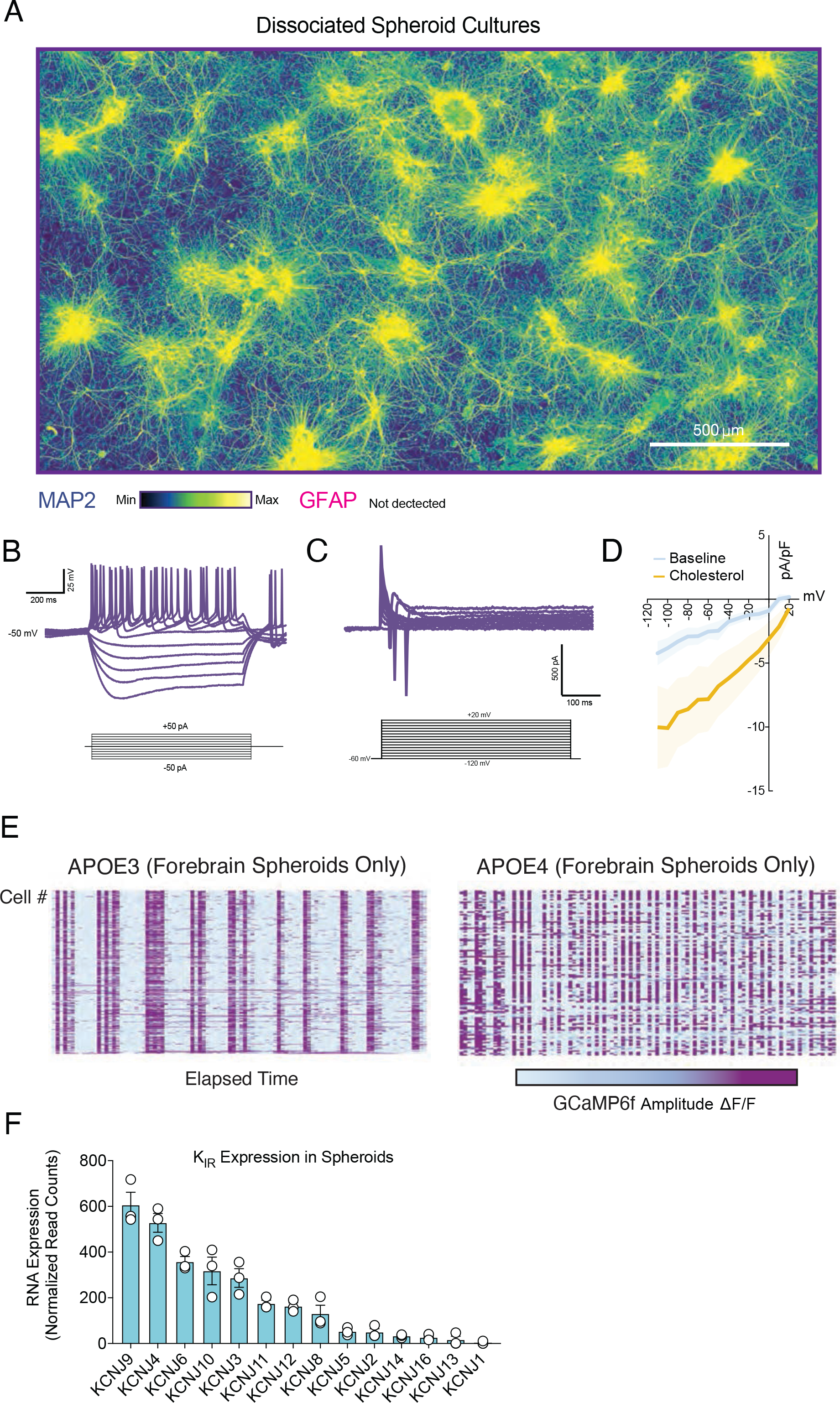
Neuronal Network Dynamics is Remodeled by APOE4. Dissociated spheroid cultures generated from APOE3 or APOE4 isogenic lines exhibit striking differences in neuronal activity patterns. (A) Large tiled image of dissociated spheroid neuronal cultures stained for MAP2 and shown in an signal intensity gradient from purple (min) to yellow (max). GFAP staining did not detect any positive cells, which is expected as astrocytes only become detectable after approximately 120 days. (B-C) Traces of whole-cell current-clamp and voltage-clamp recording shown in main figure 5G; Evoked action potentials display mature firing properties. (D) Current-voltage relationship (I-V curve) for APOE3 neurons treated with 1mM cholesterol shows potentiation of inwardly rectifying K^+^ currents. n= 6-7 cells per group. (E) Spontaneous calcium transients of APOE3 or APOE4 neurons labeled with AAV pSYN-GCaMP6f, shown in heatmaps generated from fluorescence signal time series (ΔF/F: change in fluorescence divided by baseline fluorescence). Calcium transients are shown as a raster plot. (F) RNA-seq for all Kir channels genes detected in APOE3 spheroids.

**Supplemental Figure 4.**
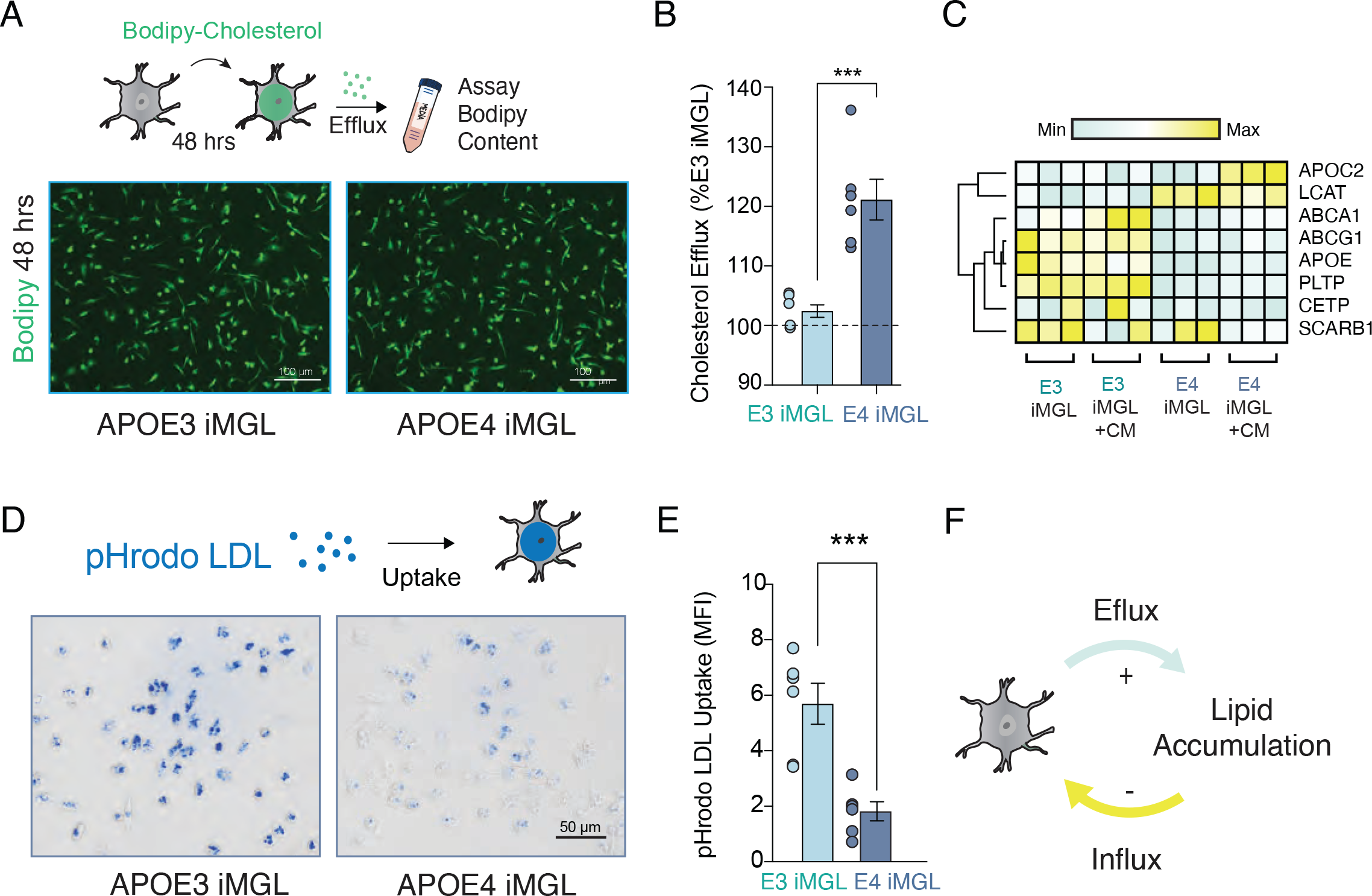
Decreased Lipid Re-Uptake by APOE4 iMGL Leads to Extracellular Accumulation. (A) APOE3 and APOE4 iMGLs were incubated with Bodipy-Cholesterol for 48hrs to saturate cholesterol uptake levels, then incubated for an additional 24 hours with phenol-free iMGL media.(B) Bodipy signal was measured from media using a plate-reader assay. Unpaired t- test, *** p-value <0.001. n= Media sampled from 6 distinct biological replicates. (C) Although we detect supernatant accumulation of APOE and cholesterol, lipid transporters are downregulated in APOE4 iMGLs. Additional genes from RNA-seq data presented in figure 3 shown as a heat map for fold-change levels and normalized as min and max per row. (D-F) pHrodo LDL uptake assay shows deficiency in lipid re-uptake, suggesting extracellular cholesterol accumulation in APOE E4 iMGLs is a result of net flux imbalance. iMGLs were exposed to pHrodo-conjugated LDL for 1 hour before imaging. Unpaired t-test, *** p-value <0.001. n= Averaged results of 3 fields of view from 6 distinct biological replicates per group.

**Supplemental Figure 6.**
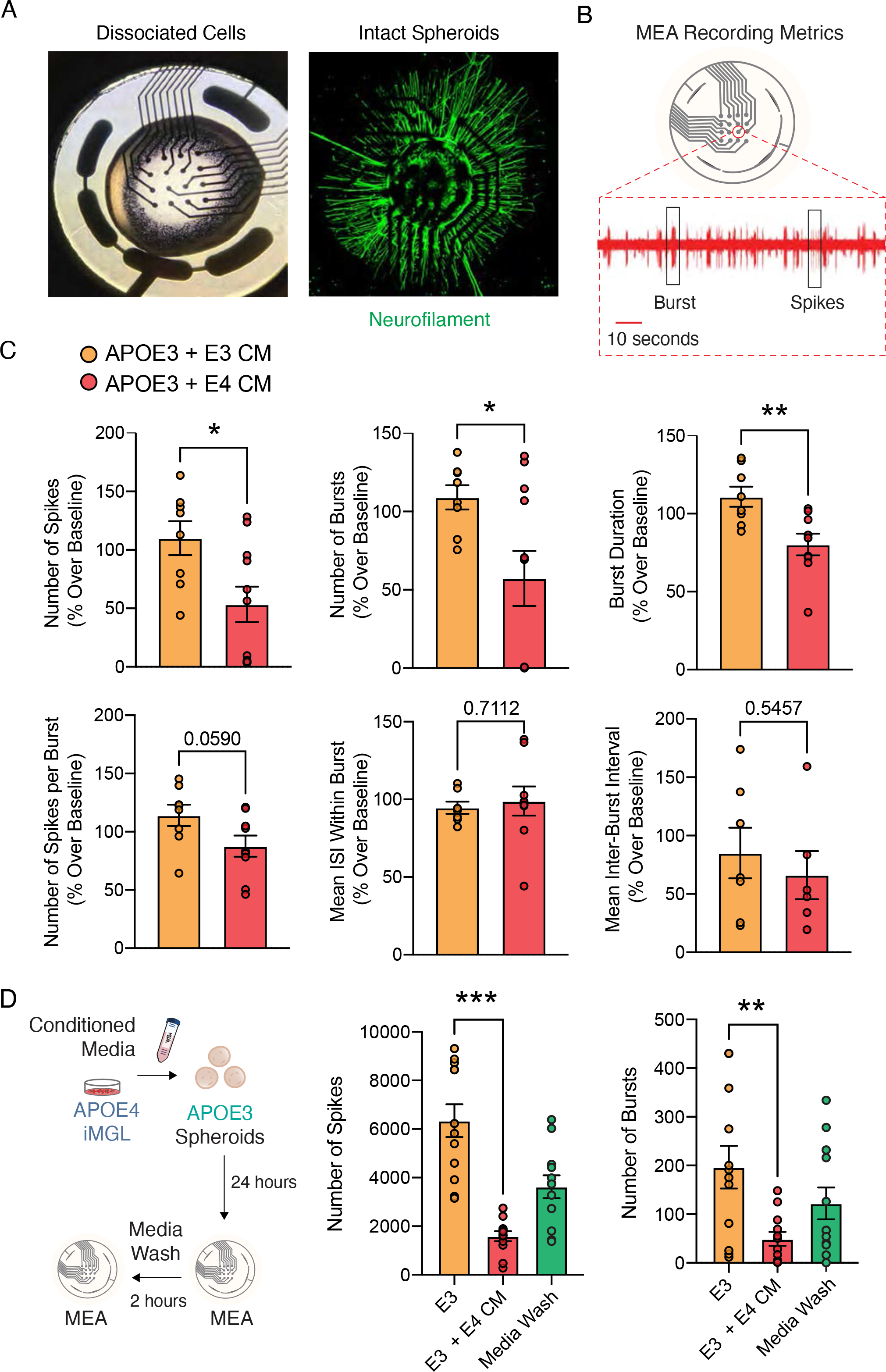
MEA Metrics for APOE3 Neurons exposed to APOE4 iMGLs Conditioned Media. (A) Dissociated cells or intact spheroids can be seeded on MEAs. (B) Spikes and bursts were measured with a MAESTRO Pro (Axion Biosystems) multielectrode array in a 48-well format with 16 electrodes following default settings. (C) Spike and Burst metrics for intact APOE3 spheroids seeded on MEA and exposed to APOE3 or APOE4 iMGLs conditioned media. Values normalized to baseline recording which was immediately before media change, and 24 hour incubation. Unpaired t-test, ** p-value <0.01; * p-value <0.05. n= Averaged values for 4 to 8 distinct wells per group. (D) APOE3 neurons exposed to APOE4 iMGL conditioned media for 24 hours shows reduced number of spikes and bursts, but 2 hours after washing out APOE4 iMGL media (Media Wash) values are partially restored. Unpaired t-test, ** p-value <0.01; *** p-value <0.001. n= Averaged values for 8 distinct wells per group.

## Victor_Supplemental Materials

**Supplemental Movie 1**. Ca^2+^ transients in iMGLs (Related to Main **Figure 1).** Baseline (1mM Caged ATP) and post-photostimulation (Uncaged ATP) for iMGLs in monoculture derived from healthy control subject. Live-imaging series with FLUO-4 AM calcium indicator pseudocolored from minimum (blue) to maximum (yellow) levels.

**Supplemental Movie 2**. Pacing Spheroids with Paddle Carbon Electrode (Related to Main **Figure 1).** Live imaging during stimulation. Increased calcium signal in AAV pSYN-GCaMP6f labeled neurons are shown with stimulus onset. The frame capturing the stimulus onset is frozen for 3 seconds to highly global increase in calcium influx.

**Supplemental Movie 3**. Isolectin IB4 Labeled iMGLs (Related to Main **Figure 4).** Isolectin IB4 Alexa Fluor 594 Conjugate preferentially labels microglia and localizes mostly in the soma (Shown in red). The motility of fine processes that are not captured by isolectin staining can be seen under bright field microscopy. 35 frames captured every 30 seconds shown at 60 fps.

**Supplemental Movie 4**. Calcium Dynamics of Neurons from APOE3 and APOE4 Spheroids in Dissociated Cultures (Related to Main **Figure 3 and Supplemental Figure4).** Neurons labeled with AAV pSYN-GCaMP6f pseudocolored in gradient from minimum (purple) to maximum (yellow). Video of 60 frames captured every 15 seconds shown at 10 fps.

**Supplemental Movie 5**. Conditioned Media from APOE4 iMGLs Disrupts Neuronal Activity of APOE3 Spheroids (Related for Main **Figure 5**). APOE3 and APOE4 iMGLs were pre-conditioned in neuronal media to allow for media carry over experiments with minimal disturbances to receiving neuronal cultures. APOE3 neurons from dissociated spheroids labeled with AAV pSYN-GCaMP6f pseudocolored in gradient from minimum (purple) to maximum (yellow) reporter intensity. Video captured 24 hours after half media switch. Video of 60 frames captured every 15 seconds shown at 10 fps.

**Supplemental Table 1**. KEGG Pathway analysis for up-regulated DEGs in APOE3 iMGLs exposed to conditioned APOE3 neuronal media (APOE3 vs CM).

**Supplemental Table 2**. KEGG Pathway analysis for up-regulated DEGs in APOE4 iMGLs exposed to conditioned APOE3 neuronal media (APOE4 vs CM).

**Supplemental Table 3**. KEGG Pathway analysis for down-regulated DEGs in APOE4 iMGLs exposed to conditioned media (APOE4 CM) in contrast to APOE3 iMGLs exposed to conditioned media (APOE3 CM) (APOE3 CM vs APOE4 CM).

**Supplemental Table 4**. KEGG Pathway analysis for up-regulated DEGs in APOE4 iMGLs exposed to conditioned media (APOE4 CM) in contrast to APOE3 iMGLs exposed to conditioned media (APOE3 CM) (APOE3 CM vs APOE4 CM).

